# Exploiting Vitamin B6 Dependency: BVL3572S Inhibits HisC and AlaA to Kill *Mycobacterium tuberculosis*

**DOI:** 10.1101/2025.10.27.684782

**Authors:** Zainab Edoo, Astrid Lenne-Delmotte, Camille Grosse, Marie Devaere, Marion Michel, Guillaume Caron, Ernesto Anoz-Carbonell, Kamel Djaout, Rosangela Frita, Cyril Gaudin, Line Hofmann, Sophie Lecher, Véronique Megalizzi, Alessia Michelotti, David Rengel, Pauline Rouan, Stephanie Slupek, Lina Tawk, Rudy Antoine, Hanna Kulyk, Glenn Dale, Christophe Guilhot, Nicolas Willand, Guy Lippens, René Wintjens, Alain Baulard

## Abstract

Tuberculosis remains the leading cause of death from a single infectious agent worldwide, and the growing prevalence of multi-drug resistant *Mycobacterium tuberculosis* (*Mtb*) underscores the urgent need for antibiotics with novel mechanisms of action. Here, we characterize BVL3572S, a hydroxamic acid-containing compound that is bactericidal and potently inhibits the growth of both extracellular and intracellular *Mtb*. Integrated transcriptomic, genetic, and biochemical analyses identified the pyridoxal phosphate (PLP)-dependent aminotransferases HisC (Rv1600) and AlaA (Rv0337c; formerly AspC) as the primary molecular targets of BVL3572S, thereby simultaneously impacting L-histidine and L-alanine biosynthesis. Spontaneous resistance mutants harbored mutations in *hisC* or *alaA*. Target engagement was further supported by overexpression studies: AlaA overexpression increased resistance in the presence of L-His whereas HisC overexpression paradoxically increased susceptibility. X-ray crystallography revealed a covalent adduct between PLP and BVL3572S within the HisC active site. The short occupancy of this adduct suggests a futile cycle that sequesters PLP. Isotopic labeling revealed widespread perturbation of amino acid biosynthesis, consistent with PLP starvation. The stepwise resistance observed upon supplementation with L-His and L-Ala together or with PLP alone suggests inhibition of multiple targets. Genome-scale CRISPRi and Tn-seq analyses additionally indicated disruptions in central metabolism, cell envelope integrity, and redox balance, possibly due to PLP depletion cascades. Consistent with its inhibition of AlaA, BVL3572S displayed strong synergy with D-cycloserine, a second-line antitubercular drug targeting D-alanine synthesis and impacting peptidoglycan synthesis, highlighting the potential of this compound in combination therapy. Collectively, our findings establish BVL3572S as a promising lead compound acting through a previously unexploited, multitarget mechanism that induces broad metabolic stress in *Mtb*, offering a novel therapeutic strategy against drug-resistant tuberculosis.

## Introduction

Tuberculosis (TB) remains a major global health threat, continuing to be a leading cause of morbidity and mortality worldwide. In 2023, an estimated 10.8 million new cases of active TB and 1.25 million deaths were reported (1). Despite the introduction of newer regimens containing bedaquiline and pretomanid, the emergence of drug-resistant TB remains a hurdle that has yet to be surmounted. In addition, prolonged treatment durations and adverse effects associated with current therapies contribute to poor patient adherence and increased rates of treatment failure. These challenges underscore the urgent need for shorter, more effective, and better-tolerated TB treatment regimens. Moreover, the rise of resistance highlights the critical need for novel antibiotics with new mechanisms of action.

One promising strategy for developing such agents involves targeting essential metabolic pathways in *Mycobacterium tuberculosis* (*Mtb*), the etiological agent of TB. The clinical success of bedaquiline, which targets the ATP synthase complex, has reinforced the potential of this approach. Among these pathways, de novo amino acid biosynthesis pathways are attractive targets due to the essential roles of amino acids in *Mtb* physiology. Amino acids serve not only as the building blocks of proteins but also as key components of the peptidoglycan, an indispensable component of the bacterial cell wall and they are involved in numerous regulatory and metabolic processes, making their biosynthetic enzymes attractive targets for antimycobacterial drug development.

In this context, the enzymes HisC and AlaA are compelling but previously unexploited antibiotic targets. HisC (Rv1600) is a histidinol-phosphate aminotransferase involved in L-histidine (L-His) biosynthesis. AlaA (Rv0337c) was originally annotated as AspC but it was recently shown to catalyze L-alanine (L-Ala) synthesis from pyruvate; we thus propose to change its name to AlaA (2). Both enzymes are pyridoxal phosphate (PLP)-dependent and are essential for the biosynthesis of their respective amino acid. They were identified as highly vulnerable drug targets in a genome-wide CRISPRi screen (3). Their dependence on electrophilic PLP renders them vulnerable to inhibition through the formation of adducts between PLP and small molecules in their active site, as has been described for inhibition of other PLP-dependent enzymes (4,5).

In an attempt to expand the chemical space available for anti-tuberculosis drug discovery, we scanned historical patents describing small molecules with reported antibacterial activity. Through this approach, we identified BVL3572S, a compound containing a hydroxamic acid moiety, with documented broad-spectrum activity against both Gram-positive and Gram-negative bacteria although its mode of action had not been characterized (6–8).

We found that BVL3572S potently inhibits *Mtb* growth and is bactericidal. Genetic and biochemical analyses identify the PLP-dependent aminotransferases HisC and AlaA as its primary targets. Beyond direct enzyme inhibition, we propose that BVL3572S also depletes PLP, broadly disrupting PLP-dependent metabolism. Finally, BVL3572S synergizes with D-cycloserine, underscoring its potential in combination therapy against drug-resistant TB.

## Results

### BVL3572S exhibits potent bactericidal activity against *Mtb*

BVL3572S displayed potent growth inhibition against *Mtb* H37Rv, with a minimal inhibitory concentration (MIC_90_) of 1.7 µM **(Fig. 1A and 1B)**. *In vitro* killing assays demonstrated that BVL3572S is bactericidal at concentrations ≥10 µM, with a 2-log decrease observed at day 7 and a 4-log decrease at day 14. Moreover, no regrowth was observed as from 30 µM, indicating the potential of BVL3572S for sterilizing activity **(Fig. 1C)**.

**Fig. 1.**
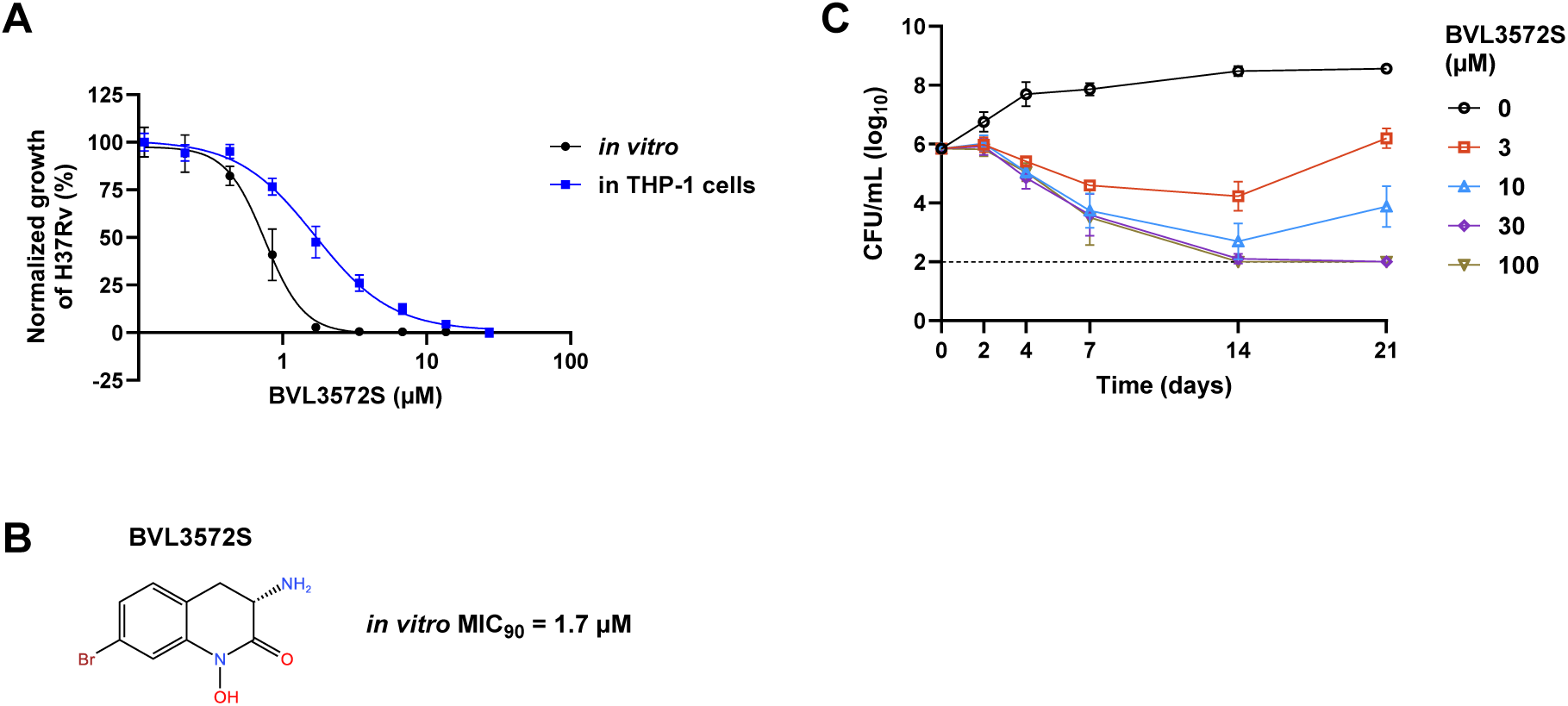
Inhibitory activity of BVL3572S against *Mtb* in axenic culture and intracellular conditions. **(A)** Dose-response analysis of BVL3572S against *Mtb* H37Rv-GFP in 7H9 medium and in differentiated THP-1 macrophages. Bacterial growth was quantified after 5 days of compound exposure by GFP fluorescence (axenic *Mtb*) or luminescence (intracellular *Mtb*). Data are means ± standard deviation (SD) of technical duplicates from one experiment representative of two to three independent biological replicates. **(B)** Chemical structure of BVL3572S and its *in vitro* MIC_90_, defined as the lowest compound concentration causing ≥90% growth inhibition. **(C)** Time-kill kinetics of *Mtb* H37Rv exposed to BVL3572S. Colony-forming units (CFUs) were enumerated on compound-free 7H10 agar at indicated time points. The dashed line indicates the limit of detection. Data points are means ± SD from three independent experiments.

BVL3572S displayed intracellular efficacy in *Mtb*-infected THP-1 macrophages, showing that BVL3572S retains activity within the host environment and can penetrate host cell membranes to reach intracellular bacteria **(Fig. 1A)**.

### Transcriptomics analysis of BVL3572S-treated *Mtb* reveals perturbation of amino acid biosynthesis

RNA-seq analysis of BVL3572S-treated *Mtb* demonstrated significant upregulation of genes involved in the L-His biosynthesis pathway, specifically *hisB* (imidazole glycerol-phosphate dehydratase), *hisH* (imidazole glycerol phosphate synthase), and *hisA* (phosphoribosylformimino-5-aminoimidazole carboxamide ribotide isomerase), as well as *alaA* (formerly *aspC*), recently characterized as encoding an alanine aminotransferase essential for *de novo* L-Ala biosynthesis (2) **(Fig. 2)**. These transcriptional changes suggested that BVL3572S may interfere with the biosynthesis of both L-His and L-Ala. Concomitantly, *whiB7*, encoding a transcriptional activator that orchestrates adaptive responses to amino acid limitation and directly regulates *alaA* transcription under alanine starvation conditions, was strongly upregulated(9). Consistent with this finding, deletion of *whiB7* decreased the MIC of BVL3572S by 4-fold compared to the wild-type (WT) strain **(Fig. 3A)**.

**Fig. 2.**
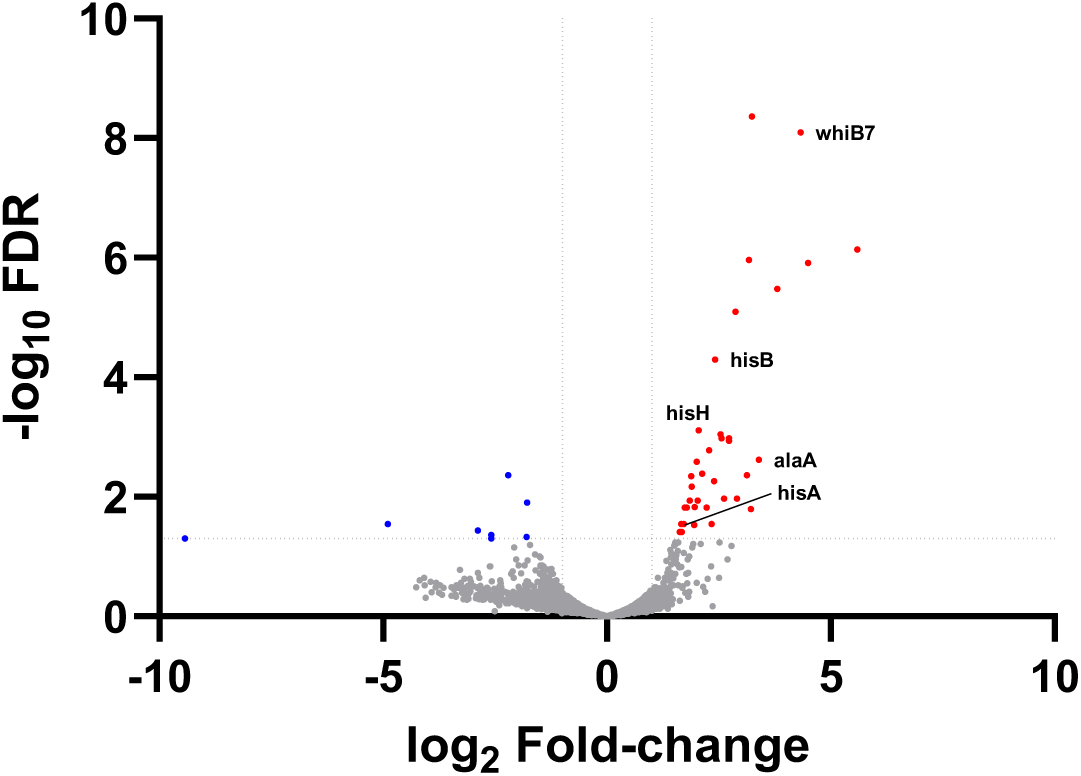
Transcriptomics profile of *Mtb* treated with BVL3572S. Volcano plot showing log_2_ fold-change (L2FC) and false discovery rate (FDR) values for each gene compared to untreated controls. Dotted lines indicate thresholds of |L2FC| > 1 and -log_10_ (FDR) > 1.3. Significantly upregulated and downregulated genes are shown in red and blue, respectively. Data represent two independent biological replicates.

**Fig. 3.**
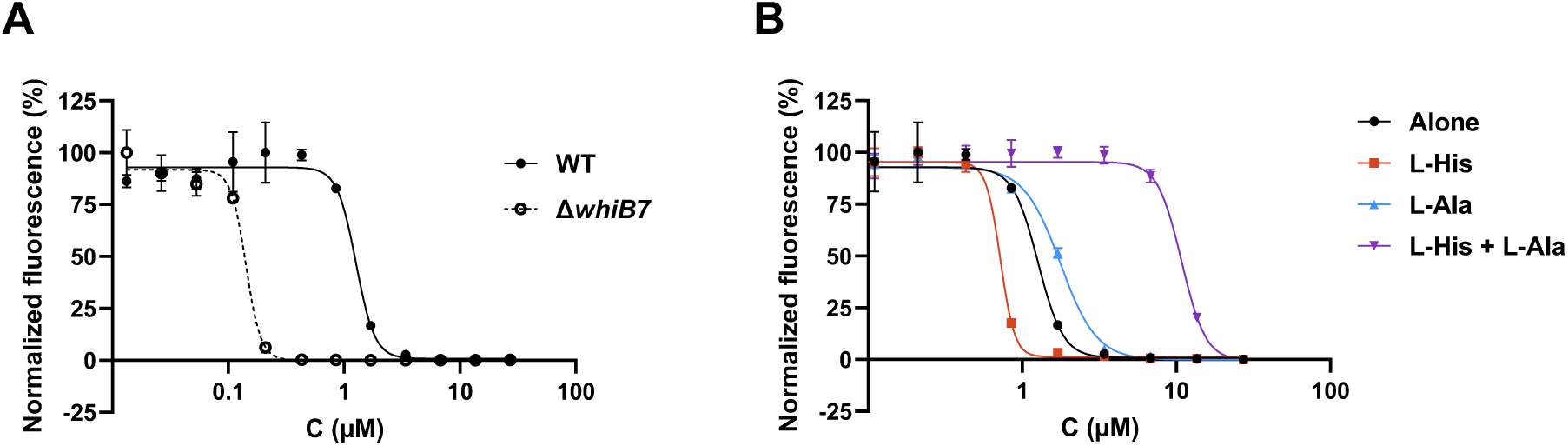
**(A)** Growth inhibitory activity of BVL3572S against *Mtb* H37Rv WT and Δ*whiB7*. **(B)** Effect of L-His and L-Ala supplementation on the growth inhibitory activity of BVL3572S against *Mtb* H37Rv. Dose-response curves represent means ± SD of two technical replicates and are representative of three biological replicates.

### Exogenous amino acid supplementation suggests dual-pathway targeting by BVL3572S

Given that our transcriptomic analysis revealed significant upregulation of genes involved in histidine and alanine biosynthesis following BVL3572S treatment, we investigated whether exogenous amino acid supplementation could rescue the predicted enzymatic deficiencies. Individual supplementation with either L-His or L-Ala failed to confer resistance to BVL3572S **(Fig. 3B).** However, simultaneous supplementation with both L-His and L-Ala conferred strong resistance to BVL3572S, suggesting that this compound might inhibit both amino acid biosynthetic pathways and demonstrating that single-pathway bypass is insufficient to overcome the growth-inhibitory effect.

Systematic amino acid rescue screening confirmed the specificity of this dual-pathway targeting: no other amino acid, tested individually or in binary combinations with L-His or L-Ala, reversed BVL3572S-induced growth arrest (data not shown). This narrow rescue profile corroborates our transcriptomic findings and strongly supports direct or indirect inhibition of L-His and L-Ala biosynthesis by BVL3572S.

### Spontaneous resistance mutations reveal HisC and AlaA as potential targets of BVL3572S

To explore the molecular targets of BVL3572S, we selected spontaneous resistant mutants under different selective pressures. Mutants selected on BVL3572S in the presence of L-His harbored an A270T substitution in AlaA, corroborating the transcriptomic evidence implicating this aminotransferase in the mechanism of action of BVL3572S **(Table 1)**. The mutants displayed conditional resistance profiles and remained viable without L-His supplementation, indicating preserved histidine biosynthesis capacity. However, they exhibited mild resistance (2-fold MIC increase) in the absence of L-His, which was enhanced 8- to 16-fold upon L-His supplementation. This phenotype suggests that the A270T substitution, located proximally to the catalytic Lys265 residue, maintains AlaA catalytic function while reducing BVL3572S binding affinity, thereby conferring partial target protection that still requires histidine pathway supplementation for maximal resistance.

**Table 1.** Spontaneous resistant mutants selected in the presence of BVL3572S and amino acids. L-Ala and L-His were supplemented at 3 mM. NA, not applicable.

| Strain | Mutant selection |  | Mutated gene | Substitution in corresponding protein | MIC<sub>90</sub> of BVL3572S (μM) |  |  |  |
| --- | --- | --- | --- | --- | --- | --- | --- | --- |
|  | Amino acid | BVL3572S (μM) |  |  | Alone | + L-His | + L-Ala | + L-His + L-Ala |
| WT | NA | NA | NA | NA | 1.7 | 0.9 | 1.7 | 27 |
| 25_1c | None | 8.5 | *alaA* | T259N | 6.8 | 6.8 | 6.8 | 27 |
| 125H1c | L-His | 4.3 | *alaA* | A270T | 3.4 | 27 | 3.4 | 13.6 |
| 5H1c | L-His | 17 | *alaA* | A270T | 3.4 | 27 | 3.4 | 27 |
| 5H3c | L-His | 17 | *alaA* | A270T | 3.4 | 13.6 | 1.7 | 13.6 |
| 25A6b | L-Ala | 8.5 | *hisC* | A147V | 0.9 | 0.9 | 27 | 13.6 |
| 5A1 | L-Ala | 17 | *hisC* | P116L | 0.9 | 0.9 | 27 | 27 |
| 5A2 | L-Ala | 17 | *hisC* | W141C | 0.9 | 0.9 | 27 | 27 |
| 10A1 | L-Ala | 34 | *hisC* | A120V | 0.9 | 0.9 | 27 | 27 |

Conversely, mutants selected in the presence of L-Ala harbored mutations in *hisC*, providing additional evidence for the role of HisC in the mode of action of BVL3572S. These mutants exhibited strict L-Ala-dependent resistance (16-fold MIC increase). Remarkably, they showed slightly enhanced susceptibility (0.5x MIC) to BVL3572S compared to WT in the absence of L-Ala supplementation, but achieved high-level resistance when L-Ala was provided. The functional impact of these substitutions, which cluster predominantly near the protein surface, is difficult to predict without direct enzyme activity measurements. However, our results suggest that these mutations might compromise catalytic efficiency while simultaneously reducing BVL3572S binding affinity.

Lastly, we selected mutants without amino acid supplementation. These exhibited modest resistance (4-fold MIC increase) and harbored a T259N substitution in AlaA. These mutants achieved high-level resistance (16-fold MIC increase) only upon simultaneous supplementation with both L-His and L-Ala, confirming obligate dual-pathway dependency.

Some mutants contained additional secondary mutations elsewhere in the genome although these are not expected to contribute to BVL3572S resistance **(Supp. Table 1)**.

### Target validation in *E. coli* confirms HisC as the primary target of BVL3572S

To validate target engagement and assess the contribution of individual pathways to BVL3572S susceptibility, we employed *E. coli* as a heterologous bacterial model. WT *E. coli* exhibited sensitivity to BVL3572S that was completely abolished by L-His supplementation, indicating that HisC of *E. coli* (HisC_Ec_) may also be a target of BVL3572S **(Supp.** Fig. 1). L-Ala addition provided no protective effect, which might be attributed to the presence of three redundant alanine aminotransferases (*alaA*, *alaC*, and *avtA*) in *E. coli*.

This specific targeting in *E. coli* provided an ideal experimental system to assess the vulnerability of *Mtb* HisC (HisC_Mtb_) independently of AlaA_Mtb_-mediated effects. To directly demonstrate HisC_Mtb_ engagement in drug activity, we complemented an *E. coli* Δ*hisC*_Ec_ auxotroph with *hisC*_Mtb_ **(Fig. 4A)**. Induction of *hisC*_Mtb_ expression by L-arabinose restored histidine prototrophy, indicating that HisC_Mtb_ functionally compensated for the absence of HisC_Ec_ **(Fig. 4A)**. The complemented strain was slightly more susceptible than the WT strain at the lowest concentration of L-arabinose tested (0.016%) **(Fig. 4B and Supp.** Fig. 2A). Stepwise induction of HisC_Mtb_ progressively increased resistance to BVL3572S, with the highest L-arabinose concentration (0.25%) yielding resistance equivalent to that conferred by L-His supplementation **(Fig. 4B and 4C)**. Susceptibility to the control antibiotic rifampicin was independent of the level of HisC_Mtb_ induction, indicating that BVL3572S resistance was due to specific target overexpression rather than improved growth of the complemented strain **(Fig. 4D)**. As additional controls, we showed that L-arabinose or the empty vector alone did not impact the susceptibility of the WT strain to BVL3572S and that increasing HisC_Mtb_ expression in the WT strain conferred resistance to BVL3572S **(Supp.** Fig. 2). These results indicate that HisC_Mtb_ expression allowed growth of *E. coli* Δ*hisC*_Ec_ in the absence of L-His supplementation and specifically conferred resistance to BVL3572S.

**Fig. 4.**
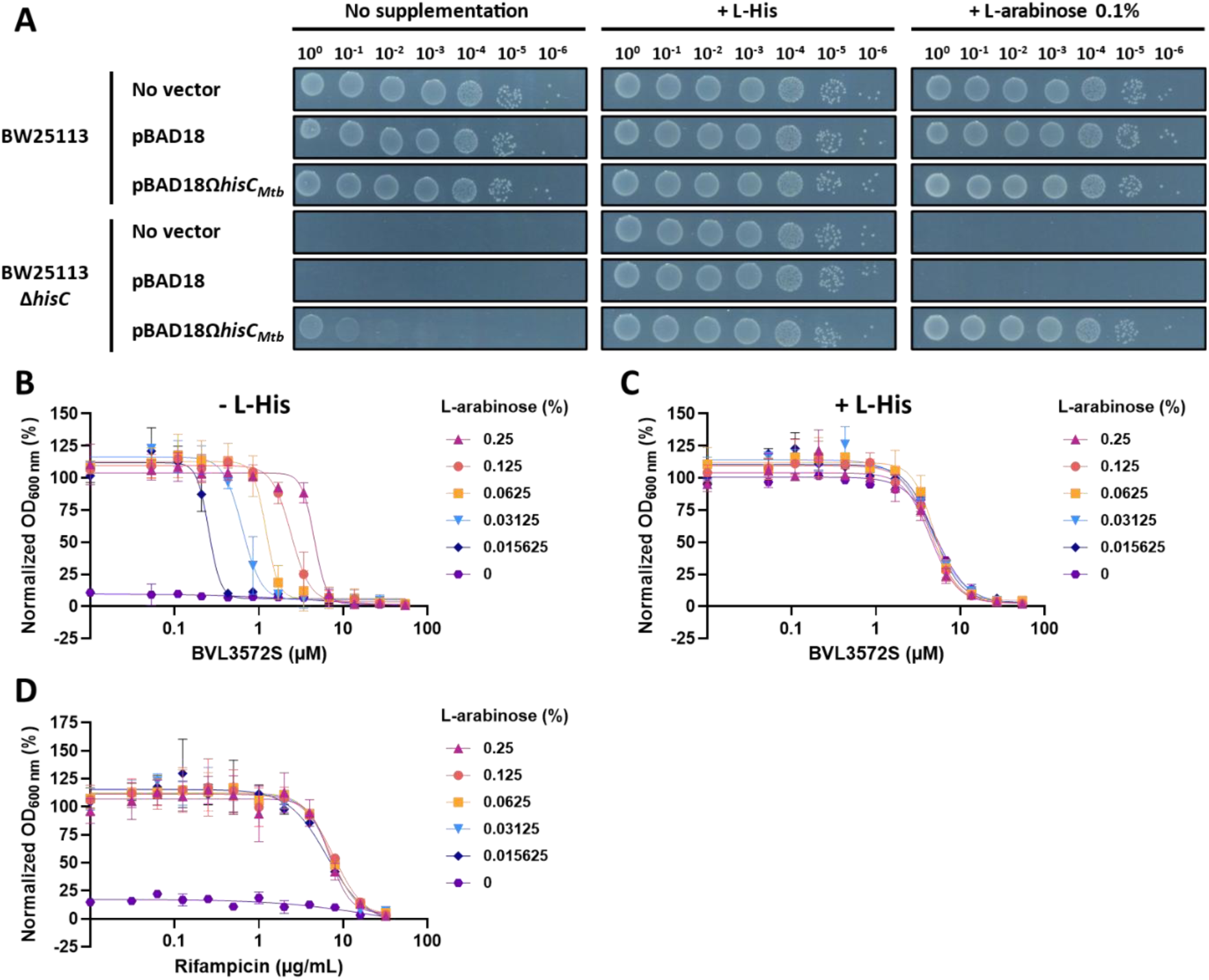
Cross-species complementation validates HisC as the target of BVL3572S. **(A)** Functional compensation of L-His auxotrophy by HisC_Mtb_ expression in *E. coli* BW25113 Δ*hisC*. Plating efficiency was assessed on M9 agar supplemented with either 80 µM L-His (permitting growth of the Δ*hisC* mutant) or 0.1% L-arabinose (for induction of *hisC*_Mtb_). **(B, C)** Growth inhibition of BW25113 Δ*hisC* (pBAD18Ω*hisC*_Mtb_) by BVL3572S in the presence of increasing L-arabinose concentrations **(B)** without or **(C)** with L-His supplementation. **(D)** Growth inhibition of BW25113 Δ*hisC* (pBAD18Ω*hisC*_Mtb_) by rifampicin in the presence of increasing L-arabinose concentrations (fitness control). Data shown are the mean ± SD of two technical replicates and are representative of three biological replicates.

This cross-species complementation experiment provides direct evidence for target engagement between BVL3572S and HisC_Mtb_ in a bacteriological context, confirming that the compound effectively inhibits the mycobacterial enzyme at concentrations relevant for antimicrobial activity. The *E. coli* model thus provides a valuable platform for dissecting individual pathway contributions and validating HisC as a tractable target for anti-tubercular drug development.

### Structural and biochemical analysis reveals HisC inhibition and PLP sequestration

To elucidate the molecular mechanism of HisC-targeting by BVL3572S, we performed biochemical and structural characterization of this interaction. Recombinant *Mtb* HisC containing a hexahistidine tag was produced in *E. coli* and purified by nickel affinity chromatography. Enzymatic assays monitoring imidazole acetol phosphate formation demonstrated concentration-dependent inhibition of purified HisC by BVL3572S **(Fig. 5A)**. High-resolution X-ray crystallography (1.7 Å) revealed formation of a covalent adduct between BVL3572S and the PLP cofactor within the HisC active site, with clear electron density defining the adduct structure **(Fig. 5B and Supplementary Table 2)**. The proposed mechanism of formation of an external aldimine linkage between BVL3572S and PLP is a nucleophilic attack by the primary amine of BVL3572S on an internal aldimine, which is initially formed between the catalytic Lys232 residue and the aldehyde group of PLP during native transamination catalysis (10). The BVL3572S-PLP adduct is stabilized by an extensive network of intermolecular interactions with conserved active site residues, including hydrogen bonds with PLP moieties as previously observed in HisC crystals containing PLP alone (11).Additionally, the hydroxamic acid moiety of BVL3572S forms hydrogen bonds with Asn176, Arg337, and Arg346 **(Fig. 5C)**.

**Fig. 5.**
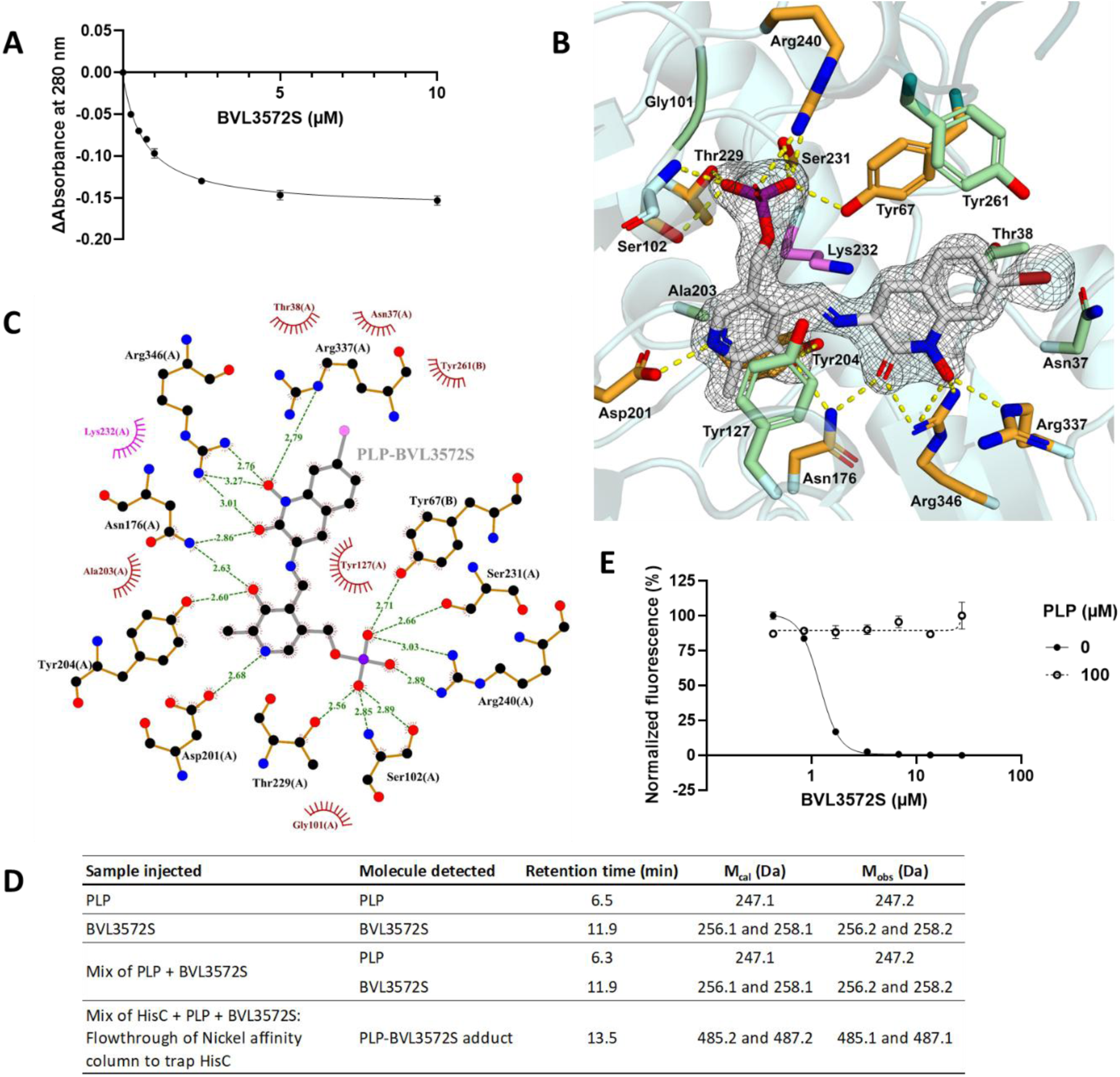
Biochemical and structural characterization of HisC_Mtb_ inhibition by BVL3572S. **(A)** Dose-dependent inhibition of purified HisC by BVL3572S monitored by measuring imidazole acetol phosphate formation. Enzymatic activity was assessed by following the enolization-dependent increase in absorbance at 280 nm. The y-axis shows decrease in absorbance at 280 nm relative to the uninhibited control. Data are means ± SD of three replicates. **(B)** Close-up view of the covalent PLP-BVL3572S adduct bound within the HisC active site. The protein is depicted as a transparent cartoon representation (chain A, pale cyan; chain B deep teal). The PLP-BVL3572S adduct and interacting HisC side chains are shown as sticks, with nitrogen, oxygen, phosphorus, and bromine atoms colored as follows: nitrogen (blue), oxygen (red), phosphorus (purple), bromine (firebrick), PLP-BVL3572S carbons (grey), hydrogen-bonding residues (bright orange), and hydrophobic contact residues (pale green). The catalytic Lys232 residue is highlighted with pink carbon atoms. Gly101 is indicated by its colored Cα atom and the Ser102 main chain is shown to display its hydrogen bond with the adduct. Hydrogen bonds are shown as yellow dotted lines, and interacting residues are labelled. The F_O_-F_C_ omit electron density map contoured at 1.0σ around the PLP-BVL3572S adduct is represented as a grey mesh. The image was generated using PyMOL v2.4.0 (Schrödinger, LLC). **(C)** 2D ligand interaction diagram of PLP-BVL3572S binding to HisC (chain A monomer). Atoms are colored as follows: oxygen (red), nitrogen (blue), carbon (black), phosphorus (violet), and bromine (pink). Hydrogen bonds are shown as green dashed lines with mean distances (Å) measured across both monomers indicated. Van der Waals contacts are represented by spoked arcs. The catalytic Lys232 residue is highlighted in pink. The image was generated using LigPlot+ (12). **(D)** LC-MS analysis of reference compounds and flowthrough fraction from Ni-NTA affinity chromatography. HisC and BVL3572S were pre-incubated at equimolar concentrations in the presence of excess PLP, then loaded onto a nickel affinity column. The flowthrough fraction was collected and analyzed by LC-MS. Individual reference compounds (PLP, BVL3572S) and their mixture without enzyme were also analyzed. M_cal_ and M_obs_ represent calculated and observed monoisotopic masses, respectively. **(E)** Effect of PLP supplementation on the BVL3572S-mediated growth inhibition of *Mtb*. Data shown are means ± SD of two technical replicates and are representative of three biological replicates.

We investigated the reversibility of adduct formation using affinity chromatography. When equimolar concentrations of HisC and BVL3572S were pre-incubated with excess PLP and applied to a nickel affinity column to retain His-tagged HisC, LC-MS analysis of the flowthrough revealed the presence of the PLP-BVL3572S adduct with the expected monoisotopic mass **(Fig. 5D)**. This demonstrates that the adduct can dissociate from the enzyme active site and suggests that BVL3572S might operate through a dual mechanism: direct enzyme inhibition via covalent adduct formation with PLP in the active site, coupled with progressive PLP depletion through iterative cycles of adduct formation and release.

To test this PLP sequestration hypothesis, we examined whether exogenous PLP could rescue BVL3572S-mediated growth inhibition in *Mtb* cultures. Strikingly, PLP supplementation completely abolished BVL3572S activity, conferring high-level resistance (MIC_90_ > 27 µM) **(Fig. 5E)**. This robust rescue effect provides compelling evidence that PLP sequestration contributes to the antimycobacterial mechanism of BVL3572S.

We performed gene overexpression studies to directly assess target engagement (**Fig. 6)**. While AlaA overexpression conferred modest resistance that was enhanced by L-His supplementation, confirming dual-pathway targeting, HisC overexpression paradoxically increased BVL3572S susceptibility. This counterintuitive result is nevertheless consistent with enhanced PLP consumption through increased formation of the PLP-BVL3572S adduct by the overexpressed enzyme, thereby accelerating intrabacterial cofactor depletion. L-Ala supplementation restored resistance in HisC-overexpressing strains by compensating for impaired alanine biosynthesis, while HisC overexpression both increased the amount of BVL3572S required for complete target engagement and possibly enhanced PLP sequestration.

**Fig. 6.**
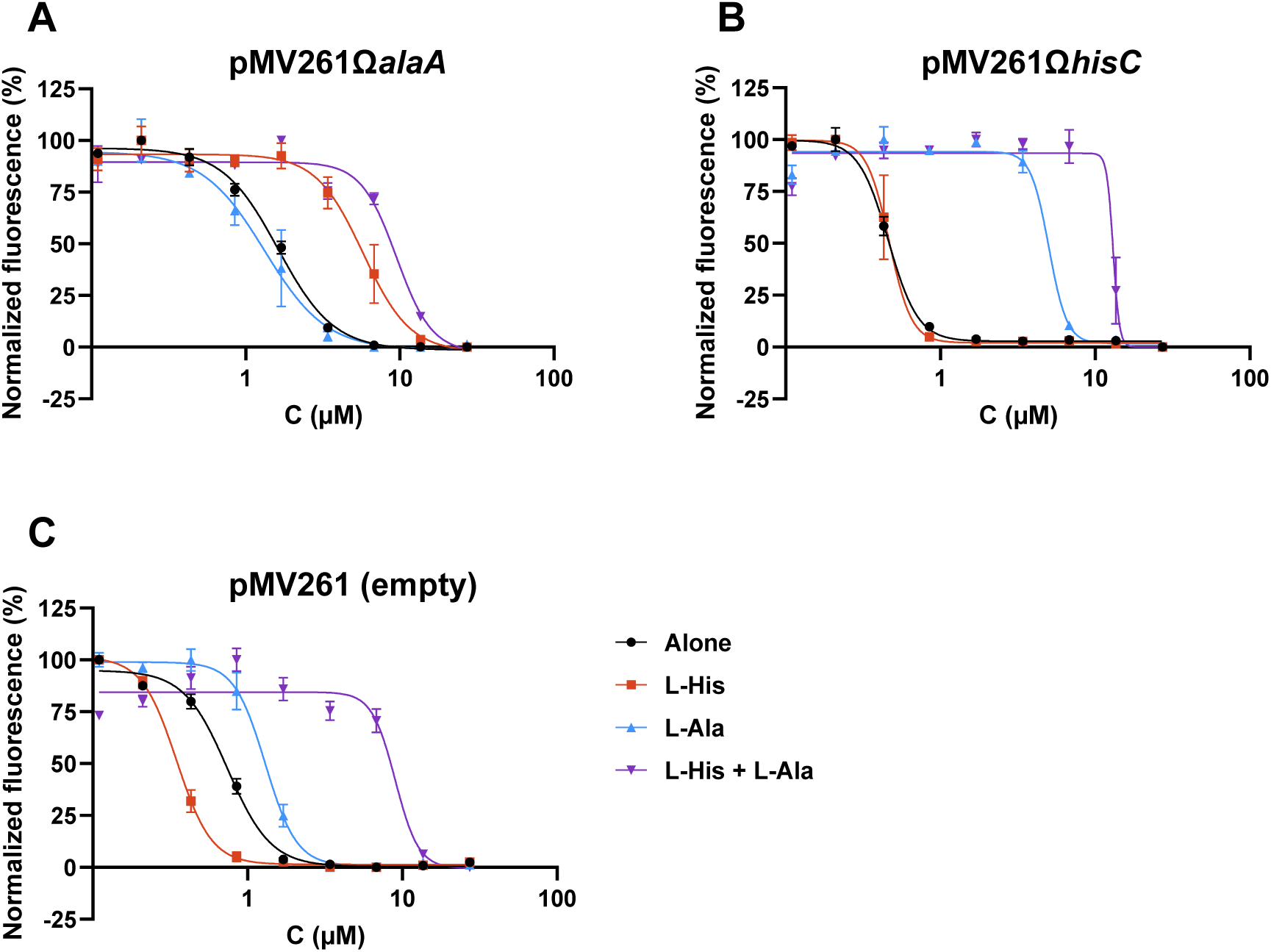
Effect of **(A)** *alaA* and **(B)** *hisC* overexpression in addition to amino acid supplementation (3 mM) on the growth inhibitory activity of BVL3572S compared to **(C)** the empty vector. Data shown are means ± SD of two technical replicates and are representative of three biological replicates.

### Exogenous amino acids do not confer resistance to intracellular BVL3572S-treated *Mtb*

We previously showed that BVL3572S inhibited the growth of intracellular *Mtb* in THP-1 cells, although to a slightly lesser extent than for extracellular bacteria **(Fig. 1A)**. To determine whether exogenous amino acids in the cellular environment could rescue intracellular *Mtb* from BVL3572S inhibition as observed in axenic culture, we supplemented infected THP-1 cells with L-His, L-Ala, or both **(Fig. 7)**. Strikingly, none of these supplementations protected intracellular *Mtb* from BVL3572S. These results suggest that while the host intracellular milieu may provide marginal protection through endogenous amino acid availability, exogenous supplementation fails to elevate intrabacterial amino acid concentrations sufficiently to overcome BVL3572S-mediated inhibition, likely due to limited uptake capacity of the cells or compartmentalization barriers in the phagosomal environment. These findings validate that endogenous amino acid salvage mechanisms are unlikely to compromise BVL3572S efficacy during infection.

**Fig. 7.**
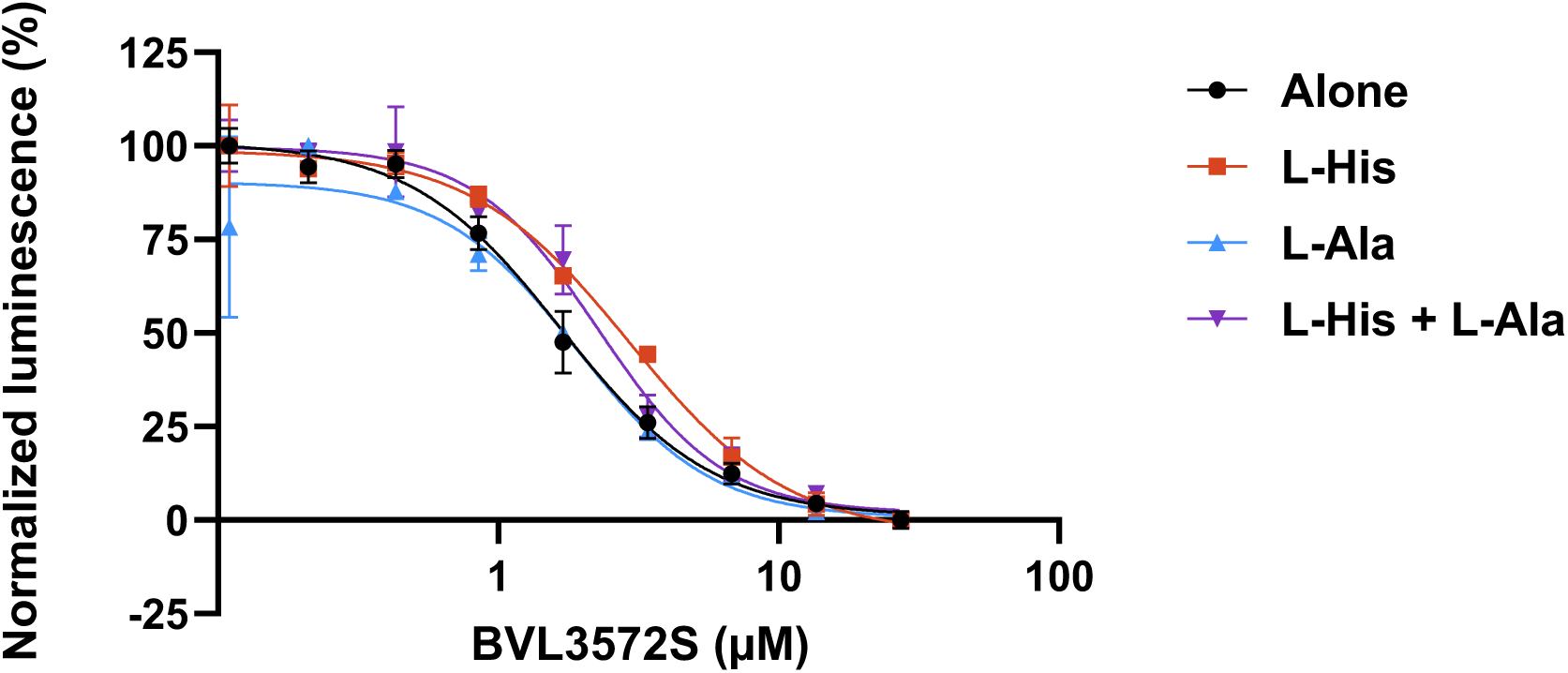
Impact of amino acid supplementation on the growth inhibitory activity of BVL3572S against intracellular *Mtb* in THP-1 cells. Data shown are means ± SD of two technical replicates and are representative of two biological replicates.

### BVL3572S treatment alters global amino acid homeostasis

Since the synthesis of most amino acids involves PLP-dependent enzymes, we hypothesized that PLP sequestration might lead to a decrease in the intrabacterial levels of several amino acids in addition to the hypothesized decrease in the intrabacterial pools of L-His and L-Ala following direct inhibition of HisC and AlaA by BVL3572S. We thus evaluated the impact of BVL3572S on the ^13^C-labeling of intrabacterial amino acids in *Mtb*. To restrain the impact of BVL3572S on bacterial viability, we limited *Mtb* treatment to less than one bacterial generation; cultures were treated for 18 h in total (2 h of initial treatment followed by 16 h of ^13^C labeling). There was a decrease in ^13^C-labeling for almost all amino acids, including histidine and alanine, consistent with transcriptomics and biochemical findings **(Fig. 8)** although no distinction between L- and D-amino acids could be made due to technical limitations. Interestingly, the only unaffected amino acid was glutamate, which is expected from the absence of involvement of PLP-dependent enzymes in its synthesis from 2-oxoglutarate. The lack of effect on glutamate labeling indicates that the impact of BVL3572S on ^13^C integration in the affected amino acids is not due to a non-specific perturbation of bacterial metabolism and viability. These results suggest that direct carbon-integrating reactions or upstream pathways involved in amino acid synthesis were impacted, possibly via PLP starvation.

**Fig. 8.**
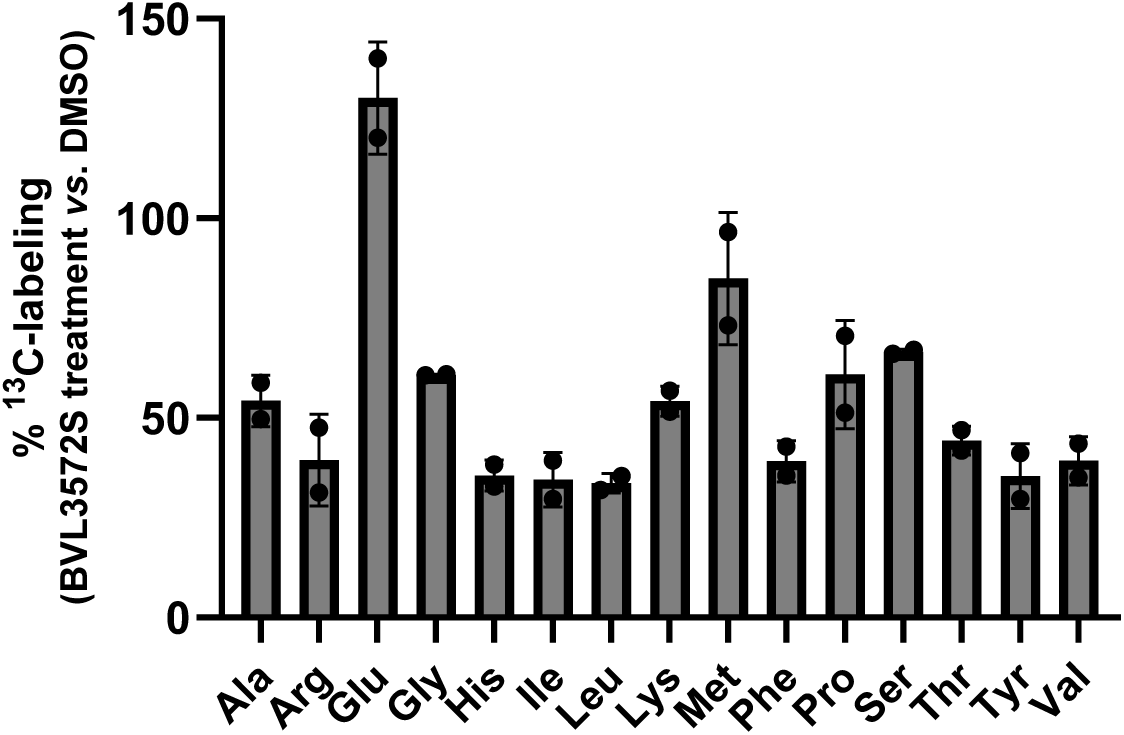
Measurement of ^13^C integration from labeled glycerol into amino acids following BVL3572S treatment compared to no treatment after OD correction. Cys, Trp, and Gln were not detected. Asp and Asn could not be distinguished because the acid treatment employed during sample preparation converted Asn to Asp. L- and D-forms of amino acids could not be distinguished. Data shown are means ± standard deviation of two biological replicates and are representative of three independent experiments.

### Genome-wide CRISPRi and Tn-seq screens reveal BVL3572S-induced metabolic imbalance

To systematically identify genetic determinants of BVL3572S susceptibility, we performed a genome-wide CRISPRi screening under four distinct metabolic conditions: without amino acid supplementation or with L-His, L-Ala, or both amino acids (**Fig. 9 and 10**). Without amino acid supplementation, *alaA* knockdown conferred increased BVL3572S susceptibility (negative fold-change), consistent with AlaA being a direct molecular target of BVL3572S. As expected, this hypersusceptibility was abolished by L-Ala supplementation **(Fig. 10A)**.

**Fig. 9.**
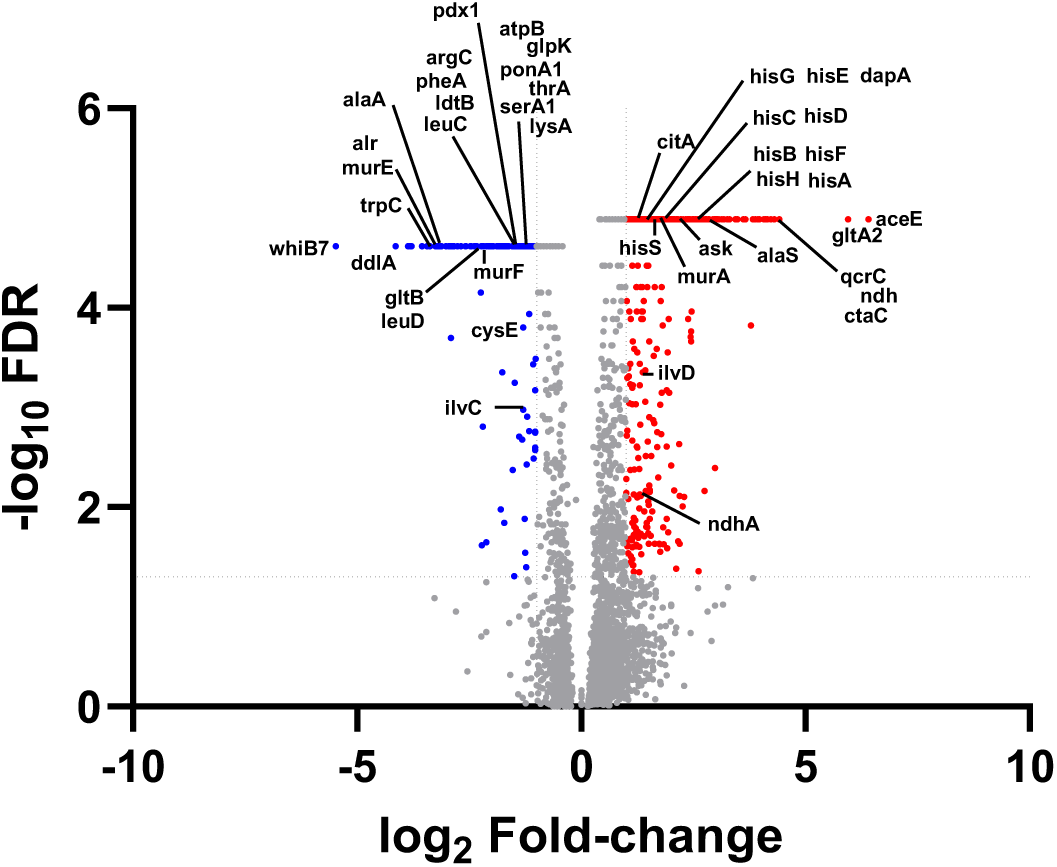
CRISPRi screen of *Mtb* treated with BVL3572S. Volcano plot showing the log_2_ fold change (L2FC) and false discovery rate (FDR) values for each gene compared to no treatment. Dotted lines represent cut-off values of |L2FC| > 1 and -log_10_ (FDR) > 1.3. Significantly depleted and enriched genes are shown in blue and red, respectively.

**Fig. 10.**
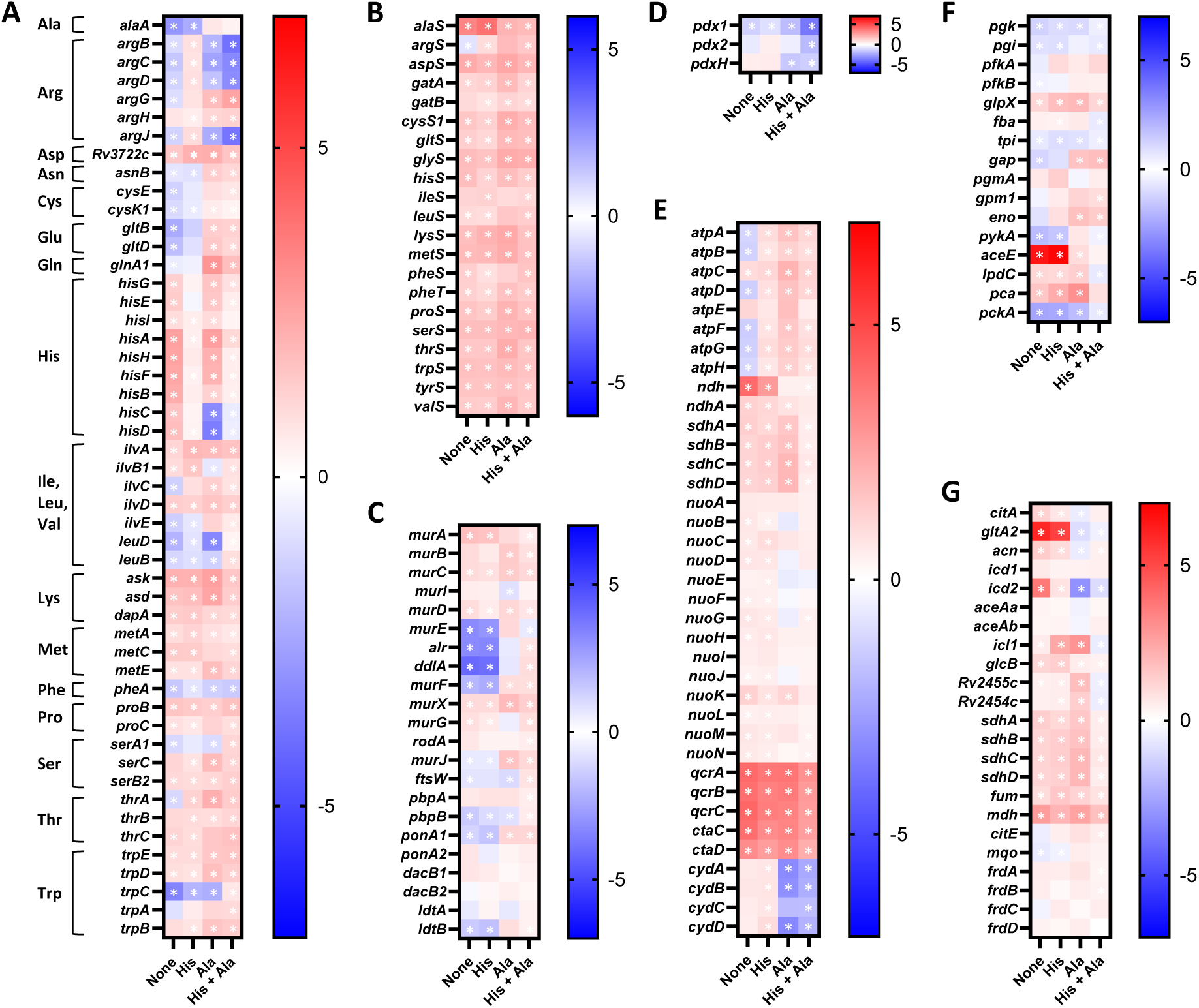
Heatmap depicting fold changes for *Mtb* genes (**A**) involved in amino acid biosynthesis, (**B**) encoding tRNA synthetases, and involved in (**C**) peptidoglycan synthesis, (**D**) PLP synthesis, (**E**) respiration, (**F**) glycolysis, and (**G**) the TCA cycle. For amino acid synthesis, only genes that were significantly enriched or depleted (|L2FC| > 1) in at least one condition are represented. Cells with a - log_10_ (FDR) > 1.3 are marked with an asterisk.

Similarly, depletion of *whiB7* increased BVL3572S susceptibility, consistent with our transcriptomic data showing strong *whiB7* upregulation upon drug treatment **(Fig. 2)** and MIC experiments demonstrating that Δ*whiB7* mutants exhibit 4-fold increased susceptibility **(Fig. 3A)**.

Interestingly, increased susceptibility to BVL3572S was also provoked by depletion of *alr* and *ddlA*, encoding an alanine racemase (conversion of L-Ala to D-Ala) and a D-Ala-D-Ala ligase (formation of a D-Ala-D-Ala dipeptide), respectively (**Fig. 9 and 10C**). Both enzymes are essential for *Mtb* growth and are involved in peptidoglycan biosynthesis. By inhibiting AlaA and reducing the intrabacterial pool of L-Ala available for racemization and integration into the peptide stems of the peptidoglycan, BVL3572S might indirectly impede peptidoglycan synthesis. This is also reflected by the increased susceptibility conferred by knockdown of genes encoding other enzymes involved in peptidoglycan synthesis, namely *murE*, *murF, ponA1,* and *ldtB*. In contrast, knockdown of early-stage peptidoglycan synthesis genes, such as *murA*, increased BVL3572S tolerance, suggesting that decreased peptidoglycan synthesis at the onset might reduce the impact of the drug on downstream integration of D-Ala in the peptidoglycan. The impact of AlaA inhibition on peptidoglycan synthesis is further reflected by the decreased vulnerability of *alr*, *ddlA*, and other peptidoglycan synthesis genes in the presence of L-Ala **(Fig. 10C)**.

Intriguingly, depletion of individual genes in the L-His biosynthesis pathway conferred tolerance to BVL3572S (**Fig. 9 and 10A**). Because L-His biosynthesis is essential for *Mtb* growth, we hypothesize that this tolerance does not result from L-His limitation *per se* but rather from depletion of L-His-biosynthetic enzymes interfering with HisC-mediated PLP sequestration. For *hisC*, the tolerance conferred by its depletion aligns with the hypothesis that HisC does not act only as a target of BVL3572S but also mediates PLP sequestration, as suggested by our aforementioned results (**Fig. 5**). For genes encoding proteins that intervene upstream of the histidine biosynthesis pathway (*hisA, hisB, hisE, hisF, and hisH*), their depletion is expected to lead to decreased production of the non-aminated substrate of HisC, imidazole acetol phosphate. PLP in the active site of HisC first interacts with the aminated substrate, L-glutamate, leading to the formation of an adduct between PLP and L-Glu (11,12). In the absence of imidazole acetol phosphate, PLP in the active site would be trapped as an adduct with L-Glu. The absence of free PLP in the active site is expected to curtail the formation of the PLP-BVL3572S adduct, thus preventing not only inhibition of HisC but also PLP sequestration by BVL3572S. Interestingly, depletion of *hisC* and *hisD* (the terminal enzyme in the L-His biosynthesis pathway) increased susceptibility to BVL3572S only in the presence of L-Ala, indicating that L-Ala supplementation shifts the inhibition burden to HisC **(Fig. 10A)**. Accordingly, addition of L-His but not L-Ala reduced the effects observed for genes of the His operon.

Knockdown of several genes encoding proteins involved in amino acid synthesis led to increased susceptibility to BVL3572S, indicating that the antibiotic has a generalized effect on amino acid synthesis either due to direct inhibition of each enzyme or more likely due to the interplay between biosynthesis pathways, which may use the same precursors **(Fig. 10A)**. Addition of L-His reduced the impact of the knockdown of genes involved in the synthesis of several amino acids, such as arginine, branched-chain amino acids, serine, threonine, and tryptophan while addition of L-Ala had a similar effect for cysteine and glutamate synthesis. Another possible reason for the widespread impact on amino acid synthesis is the suspected PLP starvation during BVL3572S treatment (**Fig. 5D and 5E**). Notably, depletion of the PLP synthase-encoding gene, *pdx1* (13), increased BVL3572S susceptibility independently of L-His or L-Ala, which is in line with the hypothesis that BVL3572S treatment leads to HisC-mediated PLP sequestration and that L-His supplementation can compensate for HisC inhibition but not PLP starvation **(Fig. 10D)**. Addition of both amino acids is thus expected to shift the burden from direct inhibition of their biosynthesis to the direct consequences of PLP starvation.

The impact of BVL3572S on amino acid synthesis and ultimately on *Mtb* protein synthesis is reflected by the tolerance conferred by reducing translation through depletion of aminoacyl-tRNA synthetase-encoding genes, which are highly vulnerable genes under native conditions (3). The increase in tolerance conferred by depletion of genes encoding aminoacyl-tRNA synthases was unchanged upon addition of either amino acid, suggesting that reducing translation might be a coping mechanism for the widespread perturbations in amino acid synthesis **(Fig. 10B)**.

BVL3572S treatment also impacted ATP production and the respiratory chain, as revealed by the increased tolerance conferred by depletion of *ndh* (type II NADH dehydrogenase), *qcrCAB* (cytochrome bc_1_ complex), and *ctaBCDE* (cytochrome aa_3_ oxidase) (**Fig. 9 and 10E**). In contrast, depletion of most genes of the ATP synthase operon (*atpA-H*) increased BVL3572S susceptibility, although addition of a single amino acid (L-His or L-Ala) was sufficient to relieve the burden on the ATP synthase (**Fig. 10E**). However, addition of L-Ala rendered genes encoding the cytochrome bd oxidase (*cydABCD*) more vulnerable, suggesting that this terminal electron acceptor, and its role in maintaining redox homeostasis, became more critical when *Mtb* was treated with BVL3572S in the presence of L-Ala.

The CRISPRi screen also uncovered a broader impact of BVL3572S on *Mtb* metabolism (**Fig. 9, 10F, and 10G**). Impact on carbon metabolism was reflected by the differential vulnerability to BVL3572S of several genes involved in glycolysis, with *aceE* depletion conferring high tolerance to BVL3572S **(Fig. 10F)**. AceE is a pyruvate dehydrogenase, catalyzing the conversion of pyruvate to acetyl-CoA, which then integrates the TCA cycle. AceE depletion is thus expected to protect against a decrease in the pool of pyruvate available for conversion to L-Ala by AlaA. Indeed, addition of L-Ala suppressed the tolerance conferred by *aceE* depletion. Perturbation of the TCA cycle was indicated by the tolerance conferred by depletion of *citA* and *gltA2* (conversion of oxaloacetate and acetyl-CoA to citrate), *mdh* (conversion of malate to oxaloacetate), and *icd2* (conversion of isocitrate to α-ketoglutarate) **(Fig. 10G)**.

We used whole-genome Tn-seq analysis as a complementary approach to study the mechanism of action of BVL3572S **(Fig. 11)**. A transposon mutant library was treated with BVL3572S at a concentration that slightly slowed growth of the library to enable the identification of mutants with altered sensitivity to the treatment. As the Tn-seq screen is inherently restricted to identification of genes that are non-essential in native conditions, it does not reveal treatment effects on *hisC* or *alaA*. Nevertheless, insertions in *whiB7* (transcriptional regulator of *alaA*) were significantly depleted in the treated transposon mutant library. Possibly as a consequence of the higher concentration of BVL3572S used for the Tn-seq analysis compared to the CRISPRi screen (27 µM *vs*. 0.4 µM, respectively), the Tn-seq analysis showed a broader tolerance mechanism in response to BVL3572S treatment. Metabolic adaptation of *Mtb* to BVL3572S was reflected by the negative fold-changes obtained for genes encoding enzymes involved in amino acid synthesis and nitrogen metabolism (*trpG*, *metC*, *glnA2*, and *glnD*), lipid metabolism (*scoB*, *mce1B*, and *cut4*), purine salvage (*purT*), phosphate acquisition (*pstC2* and *senX3*), and energy metabolism (*sdhA*, *sdhC*, *cydA*, and *cydB*). Mutants depleted in several cell envelope related genes (*drrA*, *accD1*, *lppX*, and *lprG*) were also underrepresented, suggesting that BVL3572S might impact the cell wall integrity of *Mtb* or alternatively, that these genes might be involved in permeability of the envelope to BVL3572S. For instance, *drrA* encodes the nucleotide-binding domain of an efflux pump, which might be involved in export of BVL3572S. Genes involved in combating oxidative stress (*katG*, *mutT1*, and *moeA1*) were depleted in the treatment condition, revealing a perturbation in the redox balance during BVL3572S treatment. Accordingly, there was a significant enrichment of transposon insertions in *fdhD* and *cyp123*, which encode proteins that might potentially increase the production of reactive oxygen species, suggesting that limiting perturbation of the intrabacterial redox balance could contribute to tolerance to BVL3572S. Interestingly, insertions in *glpK* were enriched, suggesting that disruption of glycerol metabolism increases BVL3572S tolerance, as observed for other antibiotics (14,15). This contrasts with the CRISPRi screen where *glpK* depletion was associated with increased vulnerability to BVL3572S. A likely explanation is that at the lower drug concentration used in the CRISPRi screen, *glpK* depletion imposes a metabolic burden on *Mtb* by impairing glycerol catabolism, reducing bacterial fitness and sensitizing *Mtb* to growth inhibition. In contrast, at the higher drug concentration used in the Tn-seq screen, loss of GlpK function would lead to metabolic stalling and a tolerant state, which is expected to reduce killing efficiency of BVL3572S, reflecting a context-dependent role of GlpK in modulating drug tolerance.

**Fig. 11.**
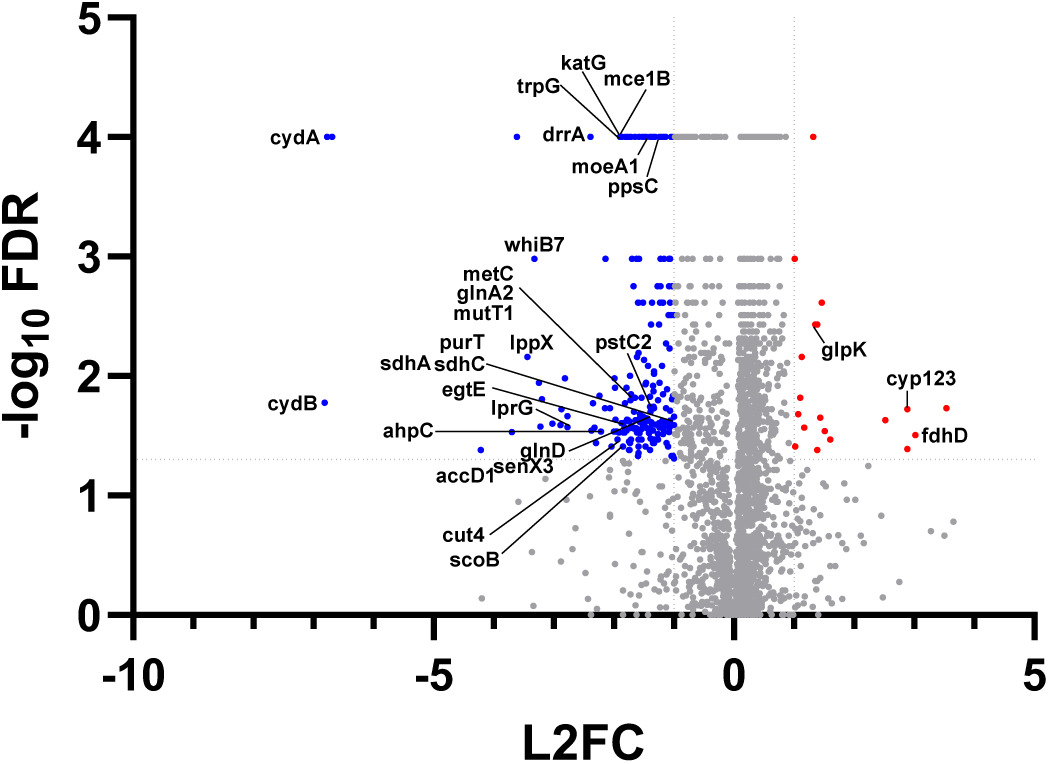
Tn-seq analysis of *Mtb* treated with BVL3572 compared to no treatment. Dotted lines represent cut-off values of |L2FC| > 2 and -log_10_ (FDR) > 1.3.

### Synergy between BVL3572S and D-cycloserine validates AlaA inhibition

Given that our CRISPRi analysis revealed perturbations in peptidoglycan synthesis pathways, we sought to evaluate the growth inhibitory activity of BVL3572S in combination with antibiotics targeting peptidoglycan biosynthesis. Combination assays demonstrated strong synergy between BVL3572S and D-cycloserine with a fractional inhibitory concentration index (FICI) of 0.15 **(Fig. 12)**. An alternative drug in MDR-TB regimens, D-cycloserine inhibits both D-alanine racemase (Alr) and D-Ala–D-Ala ligase A (DdlA) (16–19). Notably, this synergy was not observed with other peptidoglycan-targeting antibiotics, including amoxicillin, meropenem, or vancomycin, demonstrating specificity for the D-alanine biosynthetic branch rather than general peptidoglycan assembly processes **(Supp.** Fig. 3). This selective synergy supports a mechanistic model wherein BVL3572S-mediated inhibition of L-Ala biosynthesis creates a metabolic bottleneck that is further constricted by D-cycloserine-dependent downstream blockade of L-Ala to D-Ala conversion and D-Ala-D-Ala dipeptide formation, collectively impairing peptidoglycan synthesis.

**Fig. 12.**
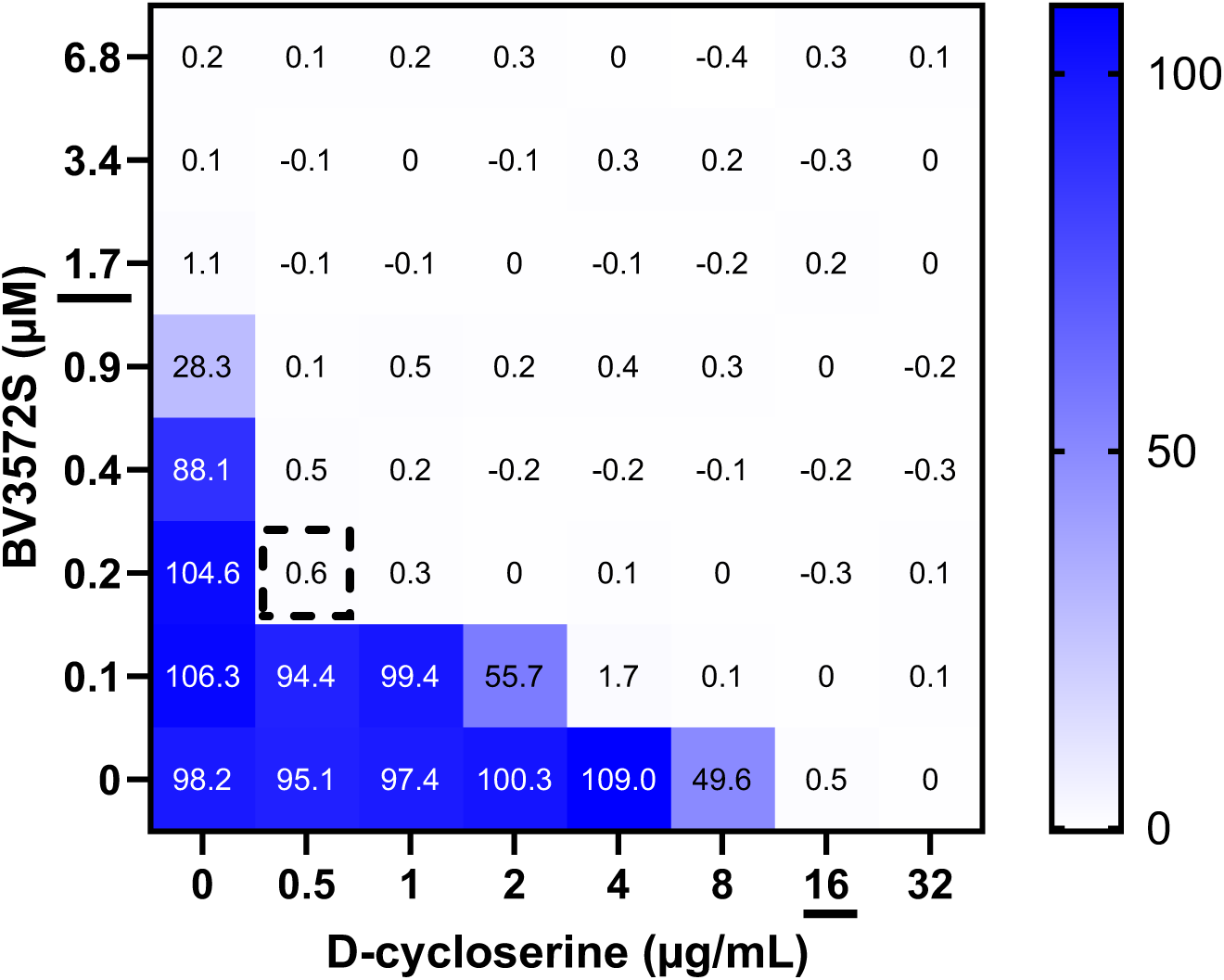
Heatmap depiction of normalized fluorescence of *Mtb* H37Rv treated with combinations of BVL3572S and D-cycloserine at the indicated concentrations. Underlined values indicate the MIC_90_ of each antibiotic, defined as the first antibiotic concentration that reduces bacterial growth by at least 90%. The dashed square shows the combination with the lowest FICI. Data represent one of three biological replicates.

Importantly, this synergy also validates the functional significance of AlaA inhibition in the mode of action of BVL3572S, demonstrating that direct disruption of alanine biosynthesis contributes substantively to the antimycobacterial mechanism of BVL3572S.

To exclude alternative mechanisms underlying this synergy, we performed comprehensive control experiments. Since Alr is also a PLP-dependent enzyme, we tested whether PLP sequestration could contribute to the synergy between BVL3572S and D-cycloserine. However, PLP supplementation conferred no protection against D-cycloserine, ruling out a possible cross-sensitization of Alr due to PLP starvation **(Supp.** Fig. 4A). Additionally, we confirmed the absence of direct cross-target inhibition by showing that overexpression of *alaA* or *hisC* had no impact on D-cycloserine susceptibility, in contrast to alr overexpression, which conferred the expected resistance **(Supp.** Fig. 4B).

In addition, evaluation of BVL3572S in combination with standard anti-TB agents revealed no antagonistic interactions **(Supp.** Fig. 3), suggesting compatibility for potential integration into existing TB regimens.

### BVL3572S analogues reveal structural determinants critical for antibacterial activity

BVL3572S analogues have previously been identified as inhibitors of human KATII for schizophrenia treatment (20), indicating that BVL3572S would require structure optimization to increase its specificity towards *Mtb*. To identify structural features essential for antibacterial activity and positions that might be amenable to modifications, we evaluated the growth inhibitory activity of a small panel of BVL3572S analogues **(Fig. 13)**. The *S*-stereoisomer was more active than the *R*-stereoisomer. Introduction of a chlorine atom at position 5 was detrimental for activity while introduction of a methoxy group was tolerated at position 7, suggesting that chemical optimization at position 7 may be feasible to increase the potency of BVL3572S and to modulate its specificity towards growth inhibition of *Mtb*. Further work is required for a comprehensive analysis of the structure-activity relationship of the BVL3572S scaffold.

**Fig. 13.**
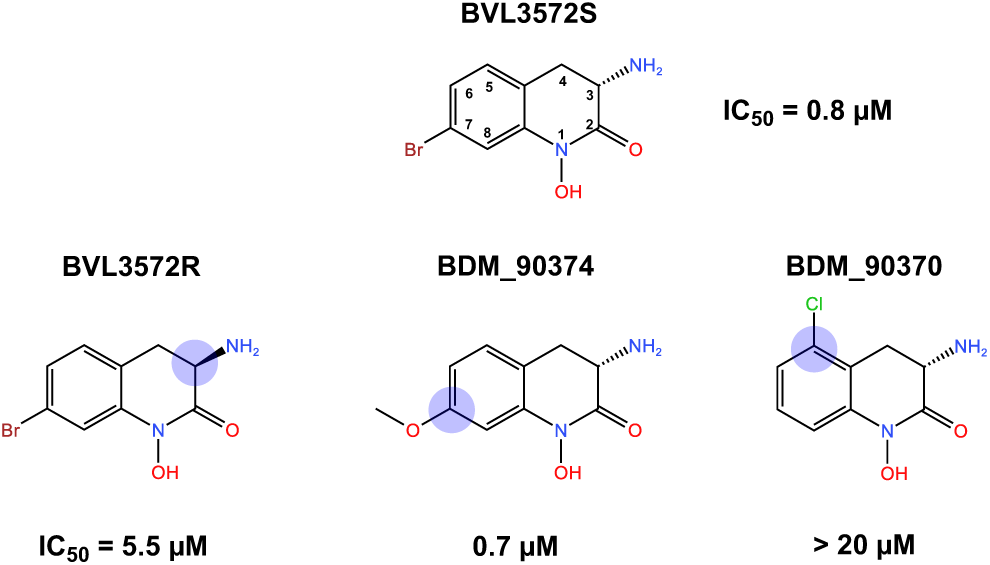
Growth inhibitory activity of BVL3572S and analogues against *Mtb* H37Rv. IC_50_ values were obtained from dose-response curves of *Mtb* grown in the presence of each compound in axenic cultures.

## Discussion

In 2023, 400,000 new cases of MDR and rifampicin-resistant TB were declared (1). Although treatment outcome for drug-resistant TB has improved in the last decade, the treatment success rate is still at 68% compared to 88% for drug-susceptible TB. Treatment of drug-resistant TB is longer, less effective, and involves drugs with toxic side-effects although the introduction of new bedaquiline-containing regimens has shortened treatment duration to 6 months (16). The increasing prevalence of drug-resistant TB exemplifies the urgent need to discover new antibiotics with novel mechanisms of action. Given the primordial role of amino acids in protein synthesis and numerous other cellular processes, amino acid biosynthesis is an interesting and as yet unexplored field for anti-TB drug development.

Our study has identified BVL3572S, a hydroxamic acid-containing compound, as a potent multitarget inhibitor of *Mtb* with a novel mechanism of action. Through extensive genetic, biochemical, and structural analyses, we demonstrated that BVL3572S exerts its bactericidal effect by simultaneously inhibiting two essential PLP-dependent aminotransferases, HisC and AlaA, thereby disrupting biosynthesis of L-His and L-Ala while possibly engendering starvation of PLP, the biologically active form of vitamin B6. This multimodal mechanism creates a synthetic lethal phenotype that extends beyond direct enzymatic inhibition to encompass systematic perturbation of amino acid metabolism and downstream effects on peptidoglycan synthesis.

A major concern for metabolic pathway inhibitors is the potential for rescue mechanisms whereby bacteria salvage missing metabolites from the host. Both L-His and L-Ala are present in human plasma at physiologically relevant concentrations and could theoretically be imported by *Mtb* during infection (21). However, it was recently shown that IFNγ-mediated signaling induces overexpression of host L-His catabolizing enzymes, resulting in significant depletion of intracellular L-His availability, forcing *Mtb* to rely on *de novo* biosynthesis, thereby validating the histidine biosynthesis pathway as a therapeutic target (22). While L-Ala can be imported from host cells (23), our results indicate that its availability is insufficient to rescue *Mtb* growth in the absence of L-His, thus exploiting the obligate dual-pathway dependency. Although simultaneous supplementation with both L-His and L-Ala partially counteracts BVL3572S activity, PLP supplementation produces complete rescue, albeit only at concentrations (100 µM) largely exceeding physiological plasma levels (8 to 165 nM) (24), indicating that endogenous PLP availability is unlikely to compromise therapeutic efficacy. Collectively, these findings suggest that metabolite salvage mechanisms are insufficient to undermine the antimycobacterial potency of BVL3572S *in vivo*.

To the best of our knowledge, our study provides evidence for the first time of PLP starvation through inhibition of PLP-dependent enzymes instead of direct inhibition of targets involved in PLP biosynthesis. The mechanism involves a futile cycle wherein BVL3572S forms a covalent adduct with PLP in the HisC active site, followed by release of the PLP-BVL3572S adduct. This iterative process is expected to progressively deplete the intrabacterial PLP pool in parallel to simultaneous BVL3572S-mediated inhibition of biosynthesis of L-His and L-Ala, creating a multitarget mode of action that extends beyond direct target inhibition. PLP sequestration is expected to amplify the effects of histidine and alanine depletion by compromising numerous PLP-dependent processes, including amino acid metabolism. Our comprehensive transcriptomics, Tn-seq, and CRISPRi analyses revealed broad metabolic dysregulation encompassing amino acid synthesis, redox imbalance, and disruption of energy-generating systems, including the electron transport chain and TCA cycle. These findings resonate with the emerging appreciation of the central role of bacterial metabolism in antibiotic efficacy and lethality (25,26).

BVL3572S impacts *Mtb* physiology at multiple interconnected levels. Direct inhibition of HisC and AlaA disrupts protein synthesis through amino acid limitation, while AlaA inhibition additionally impairs peptidoglycan synthesis by restricting L-Ala availability for downstream D-Ala formation and integration into peptidoglycan precursors. This creates an opportunity for synergistic combination with D-cycloserine, a second-line antibiotic recommended by the WHO in long-term regimens for multi-drug resistant TB (16). We demonstrated potent synergy with D-cycloserine, whereby BVL3572S-mediated L-Ala depletion synergizes with D-cycloserine-mediated downstream inhibition of L-Ala to D-Ala conversion and D-Ala-D-Ala dipeptide formation, collectively impairing peptidoglycan precursor synthesis.

BVL3572S treatment induced overexpression of WhiB7, a transcriptional regulator activated by antibiotic stress and perturbations in the biosynthesis pathways of serine, asparagine, and alanine (9,27–29). Although WhiB7 typically confers broad antibiotic resistance (27,30–32), the absence of antagonism between BVL3572S and standard TB antibiotics suggests that BVL3572S is promising as a starting chemical compound for exploration of combination regimens for MDR TB.

While metabolic pathway inhibitors have historically raised concerns about potential host toxicity, the therapeutic window for BVL3572S is potentially favorable due to a key difference between mycobacterial and human metabolism: humans lack histidine biosynthesis pathways entirely, providing selectivity for *Mtb* HisC although alanine aminotransferases are present in humans. In spite of the ubiquitous distribution of PLP-dependent enzymes across both prokaryotes and eukaryotes (33,34), multiple clinically approved drugs target PLP-dependent enzymes across diverse therapeutic areas, demonstrating that selective inhibition of PLP-dependent enzymes can be achieved with clinically acceptable safety profiles (4). In addition to D-cycloserine, other examples include vigabatrin for epilepsy (target is GABA transaminase), eflornithine for sleeping sickness (ornithine decarboxylase), and penicillamine for Wilson’s disease and rheumatoid arthritis (affecting various PLP-dependent processes) (35–37). Although the study of BVL3572S analogues as potential human KATII inhibitors for neuropsychiatric therapy showed no toxicity in preclinical studies, the compounds exhibited poor bioavailability due to O-glucuronidation at the hydroxamate moiety (20,38). While BVL3572S serves as a valuable tool compound for validating the multitarget approach of inhibiting *Mtb* growth, future development will focus on structural optimization to enhance selectivity for mycobacterial targets and improve pharmacokinetic properties.

Our study joins recent efforts exemplifying the attractiveness of metabolic pathways for TB drug discovery, including ATP synthase inhibition by bedaquiline, cytochrome bc1 inhibition by Q203, and purine synthesis inhibition targeting PurF (39–41). We validate HisC and AlaA as two additional much-needed metabolic targets for TB drug discovery, with the unique advantage of simultaneous PLP sequestration creating synthetic lethal conditions that may reduce the likelihood of resistance development in the host compared to single-target approaches. This multitarget strategy represents a promising avenue for developing next-generation antibiotics capable of addressing the urgent need for novel therapeutics against drug-resistant tuberculosis.

## Materials & Methods

### Reagents and consumables

DMSO (D8418), amoxicillin (A8523), ethambutol (E4630), erythromycin (E1300000), INH (I0500000), meropenem (M2574), PLP (82870), potassium clavulanate (33454), rifampicin (R3501), and vancomycin (V2002) were purchased from Sigma. Amikacin (TO-A003) was purchased from Euromedex. D-cycloserine (A18000) was purchased from Thermo Fisher. Bedaquiline (A12327) and pretomanid (A11451) were purchased from CliniSciences.

### Chemical synthesis of BVL3572S analogues

All reagent-grade chemicals and anhydrous solvents for synthesis, analysis and purification, were obtained from commercial suppliers and were used as received without further purification. Flash chromatography was performed using a Puriflash PF-430 with silica gel cartridges (Buchi silica 40 µm). ELSD and UV detection (254 nm) were used to collect the desired product. Reverse flash chromatography was performed using a CombiFlash® Rf200 with C18 cartridges (Buchi C18 40 µm). UV detection (215 and 254 nm) was used to collect the desired product.

1H NMR and 13C NMR spectra were recorded on a Bruker DRX-300 spectrometer. Chemical shifts (δ) are in parts per million (ppm). The ^1^H spectra were calibrated to signals from CD_2_Cl_2_ (δ 5.36 ppm), CDCl_3_ (δ 7.26 ppm) or methanol-d_4_ (δ 3.31 and 4.78 ppm), and ^13^C spectra from CD_2_Cl_2_ (δ 53.84 ppm), CDCl_3_ (δ 77.16 ppm) or methanol-d_4_ (δ 49.15 ppm). ^1^H NMR spectra are reported as following: chemical shift (ppm), multiplicity (s: singlet; brs: broad singlet; d: doublet; dd: doublet of doublet; t: triplet; td: triplet of doublet; m: multiplet), integration and coupling constants in Hertz (Hz). Proton and carbon signal assignments were established using COSY, HSQC-DEPT or HMBC spectra.

LC-MS Waters system was equipped with a 2747 sample manager, a 2695 separation module, a 2996 photodiode array detector (200-400 nm) and a Micromass ZQ2000 detec-tor (scan 100-800). XBridge C18 column (50 mm x 4.6 mm, 3.5µm, Waters) was used. The injection volume was 20 µL. A mixture of water and acetonitrile was used as mobile phase in gradient-elution. The pH of the mobile phase was adjusted with HCOOH and NH_4_OH to form a buffer solution at pH 3.8. The analysis time was 5 min (at a flow rate at 2 mL/min) or 30 min (at a flow rate at 1 mL/min). Purity (%) was determined by reversed phase HPLC, using UV detection (215 nm). Analytical analyses of each intermediate in the synthesis of BVL3572R were carried out using a SHIMADZU LC-20AD XR system equipped with an LCMS-2020 module, a DAD-MS detector with ESI ionization, and a Welch Boltimate® Core-Shell column (2.7 µm, 3.0 × 30 mm). The mobile phases consisted of 0.04% TFA in water (solvent A) and 0.02% TFA in acetonitrile (solvent B). Elution was performed with a gradient from 5% to 95% solvent B over 0.9 minutes, followed by a 0.1-minute hold at 95% solvent B, at a flow rate of 1.7 mL/min. All final compounds exhibited purities greater than 95%. All final compounds showed purity greater than 95%.

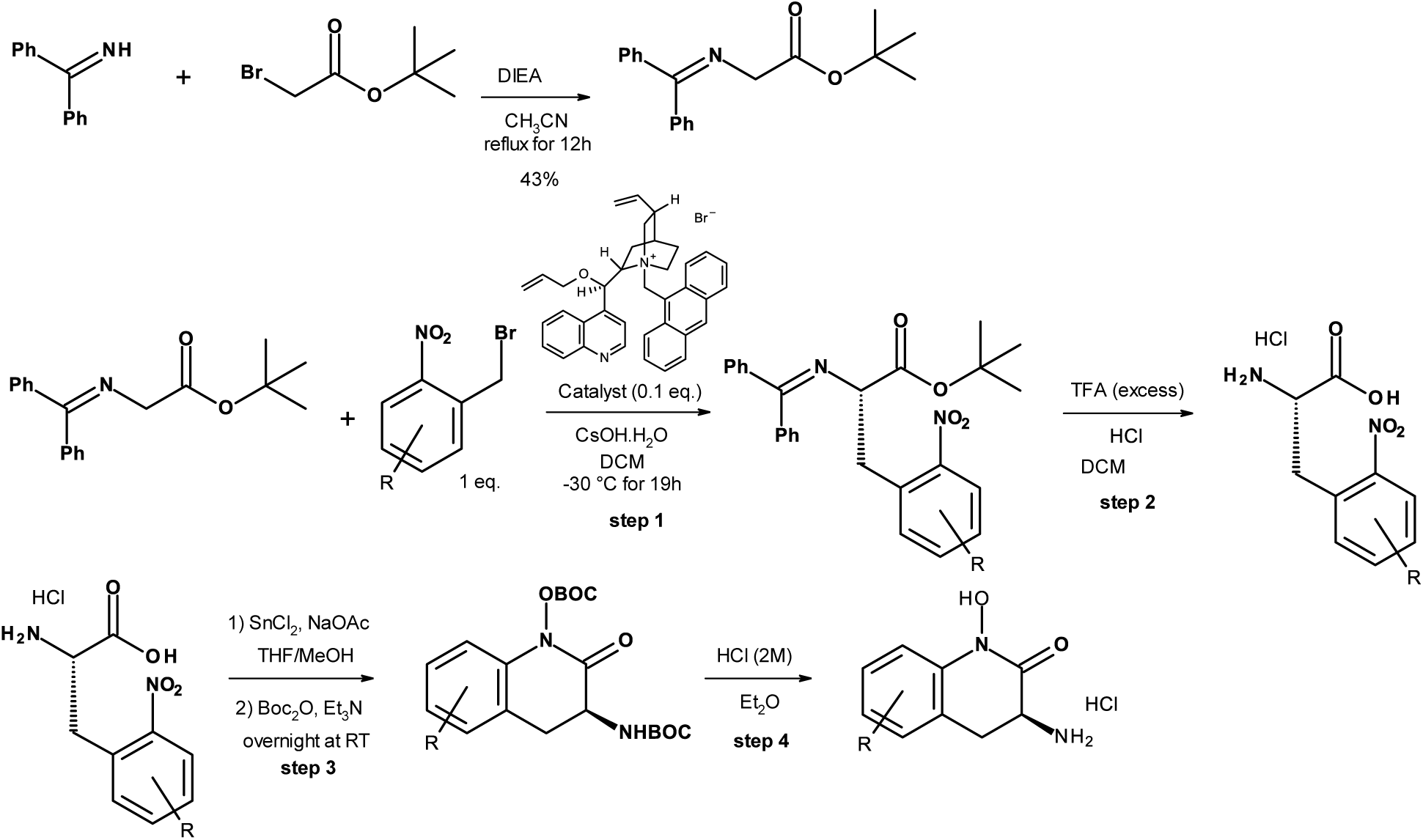

General scheme for the asymmetric synthesis of BDM_90370, BVL3572S and BDM_90374. The protocol is based on McAllister et al.(42)

### Synthesis of tert-butyl 2-(benzhydrylideneamino)acetate

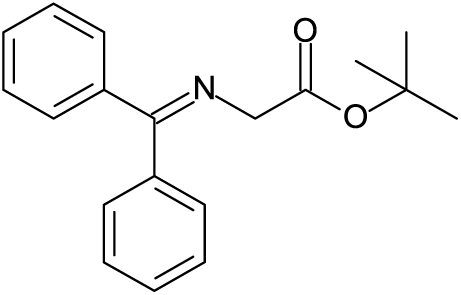

A solution of tert-butyl 2-bromoacetate (5.3 mL, 35.9 mmol) in acetonitrile (40 mL) was treated with benzophenonimine (6.0 mL, 35.9 mmol) and diisopropylethylamine (6.1 mL, 35.9 mmol), and the mixture was then heated at reflux for 12 hours. After the system had cooled to room temperature, most of the acetonitrile was removed in vacuo. The residue was partitioned between water and diethyl ether and the phases were separated. The organic layer was dried with MgSO_4_, filtered, and concentrated in vacuo until the mixture became turbid. Cooling in an ice bath provided tert-butyl 2-(benzhydrylideneamino)acetate (4.49 g, 43%) after filtration. ^1^H NMR (300 MHz, CD_2_Cl_2_) δ (ppm): 7.64 (m, 2H); 7.52-7.31 (m, 6H); 7.18 (m, 2H); 4.07 (s, 2H); 1.45 (s, 9H). LC-MS: t_r_ = 3.42 min; m/z: [M+H]^+^ 296.

### Synthesis of BDM_90374

Step 1:

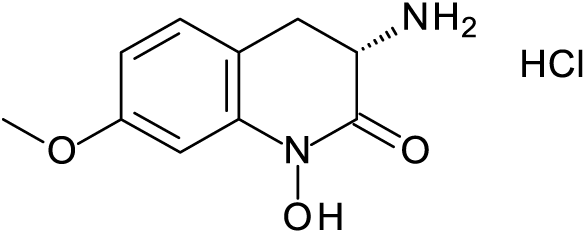

To a solution of 1-(bromomethyl)-4-methoxy-2-nitro-benzene (240 mg, 0.97 mmol, 1 equiv) in CH_2_Cl_2_ (2.8 mL) cooled to -30 °C were added tert-butyl N-(diphenylmethylidene)glycinate (330 mg, 1.12 mmol, 1.1 equiv) and 4-[(R)-allyloxy-[1-(9-anthrylmethyl)-5-vinyl-quinuclidin-1-ium-2-yl]methyl]quinoline;bromide (59 mg, 0.1 mmol, 0.1 eq). CsOH.H_2_O (219 mg, 1.46 mmol, 1.5 eq) was added and the mixture was allowed to stir for 19 h. The reaction mixture was washed with water, dried (MgSO_4_), filtered, and the solvent was removed under reduced pressure. The crude mixture was purified by flash chromatography on silica gel (Cyclohexane to 4:1 Cyclohexane:EtOAc) to give tert-butyl (2S)-2-(benzhydrylideneamino)-3-(4-methoxy-2-nitro-phenyl)propanoate (365 mg, 81%) as a yellow oil. ^1^H NMR (300 MHz, MeOD) δ (ppm): 7.52-7.26 (m, 10H); 7.11 (dd, 1H, ^3^J(H,H) = 8.61 Hz, ^4^J(H,H) = 2.81 Hz); 6.61 (d, 2H, ^3^J(H,H) = 7.07 Hz); 4.31 (dd, 1H, ^3^J(H,H) = 9.35 Hz, ^3^J (H,H) = 4.36 Hz); 3.84 (s, 3H); 3.57 (dd, 1H, ^2^J(H,H) = 13.40 Hz, ^3^J (H,H) = 3.85 Hz); 3.27 (dd, 1H, ^2^J(H,H) = 13.58 Hz, ^3^J (H,H) = 9.64 Hz); 1.46 (s, 9H). LC-MS (30 min): t_r_ = 16.50 min; m/z = [M+H]^+^ 461.

Step 2:

To a solution of tert-butyl (2S)-2-(benzhydrylideneamino)-3-(4-methoxy-2-nitro-phenyl)propanoate (315 mg, 0.62 mmol, 1 equiv) in CH_2_Cl_2_ (8.7 mL) was added TFA (8.7 mL). The reaction was allowed to stir at rt overnight. The reaction mixture was concentrated to remove TFA. The residue was partitioned between 3 M HCl and Et_2_O. The aqueous layer was washed several times with Et_2_O. The aqueous layer was then concentrated to give (2S)-2-amino-3-(4-methoxy-2-nitro-phenyl)propanoic acid;hydrochloride (181 mg, 88%) as a white solid. ^1^H NMR (300 MHz, MeOD) δ (ppm): 7.64 (d, 1H, ^4^J(H,H) = 2.77 Hz); 7.47 (dd, 1H, ^3^J(H,H) = 8.71 Hz); 7.29 (dd, 1H, ^3^J (H,H) = 8.71 Hz, ^4^J (H,H) = 2.77 Hz); 4.31 (t, 1H, ^3^J(H,H) = 7.34 Hz); 3.90 (s, 3H); 3.61 (dd, 1H, ^2^J(H,H) = 14.02 Hz, ^3^J (H,H) = 7.07 Hz); 3.34 (dd, 1H, ^2^J(H,H) = 14.02 Hz, ^3^J (H,H) = 7.86 Hz). LC-MS (30 min): t_r_ = 4.18 min, [M+H]^+^ 241

Step 3:

To a mixture of (2S)-2-amino-3-(4-methoxy-2-nitro-phenyl)propanoic acid;hydrochloride (150 mg, 0.54 mmol, 1 equiv) in THF: MeOH (16 mL: 16 mL) at 0 °C were added SnCl_2_ (514 mg, 2.71 mmol, 5 equiv) and NaOAc.3 H_2_O (738 mg, 5.42 mmol, 10 equiv). The mixture was allowed to stir, gradually warming to rt over 1.5h. Et_3_N (756 μL, 5.42 mmol, 10 equiv) and Boc_2_O (355 mg, 1.63 mmol, 3 equiv) were then added and the mixture was allowed to stir overnight at rt. The mixture was concentrated and the residue was taken up in EtOAc and H_2_O. The organic layer was washed several times with water, dried (MgSO_4_), filtered, and the solvent was removed under reduced pressure. The crude product was purified by flash silicagel chromatography eluting with 4:1 Cyclohexane:Ethyl Acetate to give [(3S)-3-(tert-butoxycarbonylamino)-7-methoxy-2-oxo-3,4-dihydroquinolin-1-yl] tert-butyl carbonate (70 mg, 32%) as a beige solid. ^1^H NMR (300 MHz, MeOD) δ (ppm): 7.21 (d, 1H, ^3^J(H,H) = 8.81 Hz); 6.70 (dd, 1H, ^3^J(H,H) = 8.33 Hz, ^4^J(H,H) = 2.52 Hz); 6.56 (m, 1H); 4.49 (t, 1H, ^3^J(H,H) = 10.81 Hz); 3.78 (s, 3H); 3.05 (d, 2H, ^3^J(H,H) = 10.39 Hz) 1.56 (s, 9H); 1.48 (s, 9H).

Step 4:

To a solution of [(3S)-3-(tert-butoxycarbonylamino)-7-methoxy-2-oxo-3,4-dihydroquinolin-1-yl] tert-butyl carbonate (65 mg, 0.16 mmol, 1 equiv) in Et_2_O (1 mL) was added 2 M HCl/Et_2_O (1 mL). The reaction mixture was stirred overnight at rt for 6h. The solid precipitate was filtered off and washed with Et_2_O to give (3S)-3-amino-1-hydroxy-7-methoxy-3,4-dihydroquinolin-2-one;hydrochloride (72 mg) as a white solid. ^1^H NMR (300 MHz, DMSO) δ (ppm): 10.92 (s, 1H); 8.80 (s, 3H); 7.22 (d, 1H, ^3^J(H,H) = 8.28 Hz); 7.41 (d, 1H, ^3^J (H,H) = 8.30 Hz); 6.82 (d, 1H, ^4^J (H,H) = 2.58 Hz); 6.65 (dd, 1H, ^3^J (H,H) = 8.18 Hz, ^4^J (H,H) = 2.58 Hz); 4.31 (dd, 1H, ^3^J(H,H) = 15.11 Hz, ^3^J(H,H) = 6.43 Hz); 3.75 (s, 3H); 3.18 (dd, 1H, ^2^J(H,H) = 14.29 Hz, ^3^J(H,H) = 6.73 Hz); 3.04 (t, 1H, ^2^J(H,H) = 14.62 Hz). ^13^C NMR (75 MHz, DMSO) δ (ppm): 161.8; 159.3; 140.0; 129.0; 111.8; 108.6; 99.7; 55.3; 48.4; 28.0. LC-MS (30 min): t_r_ : 3.88 min; m/z = [M+H]^+^ 209.

### Synthesis of BVL3572S

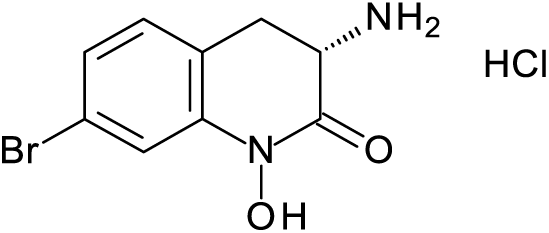

Step1:

To a solution of 4-bromo-2-(bromomethyl)-1-nitro-benzene (363 mg, 1.23 mmol, 1 equiv) in CH_2_Cl_2_ (2.8 mL) cooled to -30 °C were added tert-butyl N-(diphenylmethylidene)glycinate (400 mg, 1.35 mmol, 1.1 equiv) and 4-[(R)-allyloxy-[1-(9-anthrylmethyl)-5-vinyl-quinuclidin-1-ium-2-yl]methyl]quinoline;bromide (75 mg, 0.123 mmol, 0.1 eq). CsOH.H_2_O (277 mg, 1.84 mmol, 1.5 eq) was added and the mixture was allowed to stir for 19h. The reaction mixture was washed with water, dried (MgSO_4_), filtered, and the solvent was removed under reduced pressure. The crude mixture was purified by flash chromatography on silica gel, eluting with CH_2_Cl_2_ (isocratic) to give tert-butyl 2-(benzhydrylideneamino)-3-(4-bromo-2-nitro-phenyl)propanoate (345 mg, 55%) as a yellow oil. ^1^H NMR (300 MHz, CD_2_Cl_2_) δ (ppm): 7.99 (d, 1H, ^4^J(H,H) = 2.07 Hz); 7.59 (dd, 1H, ^3^J(H,H) = 8.28 Hz, ^4^J(H,H) = 2.07 Hz); 7.54 (dt, 2H, ^3^J(H,H) = 7.00 Hz, ^4^J(H,H) = 1.47 Hz); 7.43-7.26 (m, 7H); 6.60 (d, 2H, ^3^J(H,H) = 6.93 Hz); 4.26 (dd, 1H, ^3^J(H,H) = 9.15 Hz, ^3^J (H,H) = 4.08 Hz); 3.56 (dd, 1H, ^2^J(H,H) = 13.20 Hz, ^3^J (H,H) = 4.04 Hz); 3.35 (dd, 1H, ^2^J(H,H) = 13.47 Hz, ^3^J (H,H) = 9.30 Hz); 1.43 (s, 9H). LC-MS (5 min): t_r_ = 4.23 min; m/z: [M+H]^+^ 510.

Step 2:

To a solution of tert-butyl N-(diphenylmethylidene)-2-methyl-6-nitro-L-phenylalaninate (315 mg, 0.62 mmol, 1 equiv) in CH_2_Cl_2_ (7.4 mL) was added TFA (7.4 mL). The reaction was allowed to stir at rt overnight. The reaction mixture was concentrated to remove TFA. The residue was partitioned between 3 M HCl and Et_2_O. The aqueous layer was washed several times with Et_2_O. The aqueous layer was then concentrated to give (2S)-2-amino-3-(4-bromo-2-nitro-phenyl)propanoic acid;hydrochloride (191 mg, 95%) as a slightly yellow powder. ^1^H NMR (300 MHz, MeOD) δ (ppm): 8.29 (d, 1H, ^4^J(H,H) = 2.09 Hz); 7.89 (dd, 1H, ^3^J(H,H) = 8.36 Hz, ^4^J(H,H) = 2.09 Hz); 7.52 (d, 1H, ^3^J (H,H) = 8.21 Hz); 4.35 (t, 1H, ^3^J(H,H) = 7.46 Hz); 3.64 (dd, 1H, ^2^J(H,H) = 14.20 Hz, ^3^J(H,H) = 7.33 Hz); 3.41 (dd, 1H, ^2^J(H,H) = 13.89 Hz, ^3^J (H,H) = 7.65 Hz). LC-MS (30 min): t_r_ = 4.87; m/z = [M+H]^+^ 290.

Step 3:

To a mixture of (2S)-2-amino-3-(4-bromo-2-nitro-phenyl)propanoic acid;hydrochloride (160 mg, 0.49 mmol, 1 equiv) in THF: MeOH (14.4 mL: 14.4 mL) at 0 °C was added SnCl_2_ (466 mg, 2.46 mmol, 5 equiv) and NaOAc.3 H_2_O (669 mg, 4.91 mmol, 10 equiv). The mixture was allowed to stir, gradually warming to rt over 1.5h. Et_3_N (685 μL, 4.91 mmol, 10 equiv) and Boc_2_O (322 mg, 1.47 mmol, 3 equiv) were then added and the mixture was allowed to stir overnight at rt. The mixture was concentrated and the residue was taken up in EtOAc and H_2_O. The organic layer was washed several times with water, dried (MgSO_4_), filtered, and the solvent was removed under reduced pressure. The crude product was purified by flash silicagel chromatography eluting with 4:1 Cyclohexane:Ethyl Acetate to give [(3S)-7-bromo-3-(tert-butoxycarbonylamino)-2-oxo-3,4-dihydroquinolin-1-yl] tert-butyl carbonate (122 mg, 54%) as a white solid. ^1^H NMR (300 MHz, MeOD) δ (ppm): 7.29 (dd, 1H, ^3^J(H,H) = 8.13 Hz, ^4^J(H,H) = 1.85 Hz); 7.24 (d, 1H, ^3^J(H,H) = 8.34 Hz); 7.16 (m, 1H); 4.52 (m, 1H); 3.11 (m, 2H); 1.57 (s, 9H); 1.48 (s, 9H); 1.24 (dd, 1H, ^2^J (H,H) = 14.71 Hz, ^3^J(H,H) = 8.15 Hz).

Step 4:

To a solution of [(3S)-7-bromo-3-(tert-butoxycarbonylamino)-2-oxo-3,4-dihydroquinolin-1-yl] tert-butyl carbonate (106 mg, 0.23 mmol, 1 equiv) in Et_2_O (1.5 mL) was added 2 M HCl/Et_2_O (1.5 mL). The reaction mixture was stirred overnight at rt for 6h. The solid precipitate was filtered off and washed with Et_2_O to give (3S)-3-amino-7-bromo-1-hydroxy-3,4-dihydroquinolin-2-one;hydrochloride (48 mg, 70%) as a white solid. ^1^H NMR (300 MHz, DMSO) δ (ppm): 11.07 (s, 1H); 8.81 (s, 3H); 7.37 (d, 1H, ^3^J(H,H) = 1.52 Hz); 7.28 (m, 2H); 4.40 (dd, 1H, ^3^J (H,H) = 13.96 Hz, ^3^J(H,H) = 6.48 Hz); 3.24 (dd, 1H, ^2^J(H,H) = 15.28 Hz, ^3^J(H,H) = 6.63 Hz); 3.09 (t, 1H, ^2^J(H,H) = 15.08 Hz). ^13^C NMR (75 MHz, DMSO) δ (ppm): 161.8; 140.4; 130.1; 126.1; 120.7; 119.3; 115.6; 47.8; 28.3. LC-MS (30 min): t_r_ = 5.20 min; m/z = [M+H]^+^ 259.

### Synthesis of BDM_90370

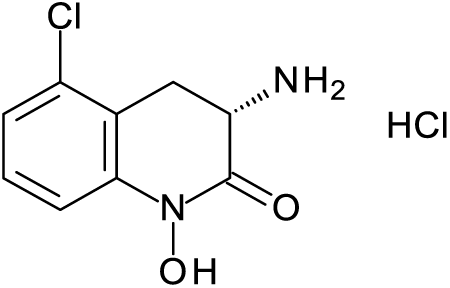

Step 1:

To a solution of 2-(bromomethyl)-1-methyl-3-nitrobenzene (308 mg, 1.23 mmol, 1 equiv) in CH_2_Cl_2_ (2.8 mL) cooled to -30 °C were added tert-butyl N-(diphenylmethylidene)glycinate (400 mg, 1.35 mmol, 1.1 equiv) and 4-[(R)-allyloxy-[1-(9-anthrylmethyl)-5-vinyl-quinuclidin-1-ium-2-yl]methyl]quinoline;bromide (75 mg, 0.123 mmol, 0.1 eq). CsOH.H_2_O (277 mg, 1.84 mmol, 1.5 eq) was added and the mixture was allowed to stir for 19 h. The reaction mixture was washed with water, dried (MgSO_4_), filtered, and the solvent was removed under reduced pressure. The crude mixture was purified by flash chromatography on silica gel, eluting with CH_2_Cl_2_ (isocratic) to give tert-butyl (2S)-2-(benzhydrylideneamino)-3-(2-chloro-6-nitro-phenyl)propanoate (430 mg, 75%). ^1^H NMR (300 MHz, CD_2_Cl_2_) δ (ppm): 7.60 (dd, 1H, 3J(H,H) = 8.09 Hz, 4J(H,H) = 1.34 Hz); 7.55-7.26 (m, 10H); 6.60 (d, 2H, 3J(H,H) = 6.93 Hz); 4.27 (dd, 1H, ^3^J(H,H) = 10.09 Hz, ^3^J (H,H) = 3.94 Hz); 3.82 (dd, 1H, ^2^J(H,H) = 13.23 Hz, ^3^J (H,H) = 10.10 Hz); 3.60 (dd, 1H, ^2^J(H,H) = 13.58 Hz, ^3^J (H,H) = 3.92 Hz); 1.43 (s, 9H). LC-MS (30 min): tr = 17.12 min; m/z: [M+H]^+^ 465.

Step 2:

To a solution of tert-butyl N-(diphenylmethylidene)-2-methyl-6-nitro-L-phenylalaninate (Fraction 1) (150 mg, 0.32 mmol, 1 equiv) in CH_2_Cl_2_ (4.3 mL) was added TFA (4.3 mL). The reaction was allowed to stir at rt for 5 min. The reaction mixture was concentrated to remove TFA. The residue was partitioned between 3 M HCl and Et_2_O. The aqueous layer was washed several times with Et_2_O. The aqueous layer was then concentrated to give (2S)-2-amino-3-(2-chloro-6-nitro-phenyl)propanoic acid;hydrochloride (71 mg, 77%) as brown solid.^1^H NMR (300 MHz, MeOD) δ (ppm): 7.99 (dd, 1H, ^3^J(H,H) = 8.14 Hz, ^4^J(H,H) = 1.32 Hz); 7.84 (dd, 1H, ^3^J(H,H) = 8.14 Hz, ^4^J(H,H) = 1.32 Hz); 7.57 (t, 1H, ^3^J (H,H) = 8.15 Hz); 4.35 (dd, 1H, ^3^J(H,H) = 9.10 Hz, ^3^J (H,H) = 8.14 Hz); 3.78 (dd, 1H, ^2^J(H,H) = 13.98 Hz, ^3^J(H,H) = 9.10 Hz); 3.49 (dd, 1H, ^2^J(H,H) = 14.07 Hz, ^3^J (H,H) = 7.28 Hz). LC-MS: t_r_ = 1.30 min; m/z: [M+H]^+^ 245

Step 3:

To a mixture of (2S)-2-amino-3-(2-chloro-6-nitro-phenyl)propanoic acid;hydrochloride (97 mg, 0.34 mmol, 1 equiv) and methyl (2S)-2-amino-3-(2-chloro-6-nitro-phenyl)propanoate;hydrochloride (3 mg, 0.01 mmol, 0.03 eq) in THF: MeOH (10 mL:10 mL) at 0 °C was added SnCl_2_ (327 mg, 1.73 mmol, 5 equiv) and NaOAc·3 H_2_O (470 mg, 3.45 mmol, 10 equiv). The mixture was allowed to stir, gradually warming to rt over 1.5h. Et_3_N (481 μL, 3.45 mmol, 10 equiv) and Boc_2_O (226 mg, 1.04 mmol, 3 equiv) were then added and the mixture was allowed to stir overnight at rt. The mixture was concentrated and the residue was taken up in EtOAc and H_2_O. The organic layer was washed several times with water, dried (MgSO_4_), filtered, and the solvent was removed under reduced pressure. The crude product was purified by flash silicagel chromatography eluting with 4:1 Cyclohexane:Ethyl Acetate to give [(3S)-3-(tert-butoxycarbonylamino)-5-chloro-2-oxo-3,4-dihydroquinolin-1-yl] tert-butyl carbonate (90 mg, 63 %) as a translucent oil. ^1^H NMR (300 MHz, CD_2_Cl_2_) δ (ppm): 7.25 (td, 1H, ^3^J(H,H) = 8.08 Hz, ^4^J(H,H) = 0.97 Hz); 7.20 (td, 1H, ^3^J(H,H) = 8.18 Hz, ^4^J(H,H) = 1.40 Hz); 6.97 (m, 1H); 5.42 (s, 1H); 4.54 (m,1H); 3.75 (dd, 1H, ^2^J(H,H) = 15.78 Hz, ^3^J (H,H) = 6.25 Hz); 2.82 (dd, 1H, ^2^J (H,H) = 15.06 Hz); 1.54 (s, 9H); 1.47 (s, 9H).

Step 4:

To a solution of [(3S)-3-(tert-butoxycarbonylamino)-5-chloro-2-oxo-3,4-dihydroquinolin-1-yl] tert-butyl carbonate (63 mg, 0.15 mmol, 1 equiv) in Et_2_O (1 mL) was added 2 M HCl/Et_2_O (1 mL). The reaction mixture was stirred overnight at rt for 6h. The solid precipitate was filtered off and washed with Et_2_O to give (3S)-3-amino-5-chloro-1-hydroxy-3,4-dihydroquinolin-2-one;hydrochloride (15 mg, 50%) as a slightly puprle solid. ^1^H NMR (300 MHz, MeOD) δ (ppm): 7.40-7.33 (m, 2H); 7.27-7.21 (m, 1H); 4.40 (dd, 1H, 3J(H,H) = 14.43 Hz, 3J (H,H) = 6.62 Hz); 3.72 (dd, 1H, 2J(H,H) = 15.57 Hz, 3J (H,H) = 6.62 Hz); 3.00 (t, 1H, 2J (H,H) = 15.62 Hz).^13^C NMR (75 MHz, MeOD) δ (ppm): 162.7; 141.6; 134.0; 130.6; 126.2; 119.0; 113.7; 49.5; 27.7. LC-MS (30 min): tr = 4.15 min; m/z: [M+H]^+^ 213.

### Synthesis of BVL3572R

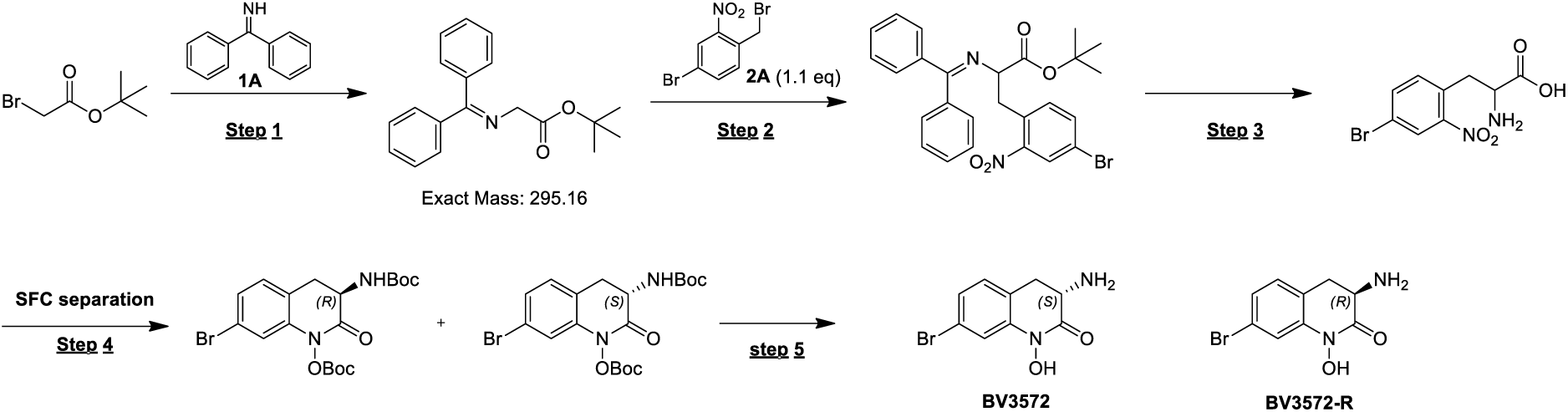

Step 1:

To a solution of tert-butyl 2-bromoacetate (24.7 g, 126 mmol, 18.7 mL, 1.15 eq) and benzophenonimine (20.0 g, 110 mmol, 18.5 mL, 1.00 eq) in ACN (240 mL) was added DIEA (16.4 g, 126 mmol, 22.1 mL, 1.15 eq). The mixture was stirred at 80 °C for 12h. The reaction mixture was then quenched by addition of water (100 mL) at 25 °C and extracted with EtOAc (2 x 100 mL). The combined organic layers were dried over Na_2_SO_4_, filtered and concentrated under reduced pressure. The crude product was triturated with petroleum ether: EtOAc = 100: 5 at 25 °C for 12 h. tert-butyl 2-((diphenylmethylene)amino)acetate (25.0 g, 84.6 mmol, 76.7% yield) was obtained as a white solid. LCMS (1 min): tr = 0.487 min; m/z = 296 [M+H]^+^.

Step 2:

To a solution of tert-butyl 2-((diphenylmethylene)amino)acetate (10.0 g, 34.0 mmol, 1.10 eq) in CH_2_Cl_2_ (20.0 mL) cooled to -30 °C, were added 4-bromo-1-(bromomethyl)-2-nitrobenzene (9.15 g, 30.9 mmol, 1.00 eq) and O-Allyl-N-(9-anthracenylmethyl)cinchonidinium bromide (1.88 g, 3.10 mmol, 0.10 eq). CsOH.H2O (7.80 g, 46.4 mmol, 1.50 eq) was added and the mixture was allowed to stir for 31 h. After this time, the reaction mixture was quenched by addition of water (100 mL) at 20 °C, and then extracted with CH_2_Cl_2_ (2 x 100 mL). The combined organic layers were dried over Na_2_SO_4_, filtered and concentrated under reduced pressure. The residue was purified by flash silica gel chromatography (ISCO®; 120 g Se pa Flash® Silica Flash Column, Eluent of 0 - 9% EtOAc/Petroleum ether gradient @ 100 mL/min). TLC (Petroleum ether: EtOAc = 10:1, Rf/p1 = 0.33), tert-butyl 3-(4-bromo-2-nitrophenyl)-2- ((diphenylmethylene)amino)propanoate (8.90 g, 17.4 mmol, 56.4% yield) was obtained as a yellow gum. LCMS (1 min): tr = 0.649 min, m/z = 510 [M+H] +.

Step 3:

To a solution of tert-butyl 3-(4-bromo-2-nitrophenyl)-2- ((diphenylmethylene)amino)propanoate (5.00 g, 9.82 mmol, 1.00 eq) in CH_2_Cl_2_ (50.0 mL) was added TFA (75.1 g, 658 mmol, 48.7 mL, 67.1 eq) and the mixture was stirred at 30 °C for 12 h. The mixture was concentrated under reduced pressure; to the residue, MeOH (10.0 mL) and HCl (3 M x 40 mL) were added and the mixture was stirred at 25 °C for 2 h. After this time, the mixture was concentrated and extracted with EtOAc (2 x 50.0 mL). The aqueous phase was filtered and concentrated under reduced pressure to give 2-amino-3-(4-bromo-2-nitrophenyl)propanoic acid (3.50 g, 12.1 mmol, 61.6% yield) as a white solid. LCMS (1 min): tr = 0.387 min, m/z = 290 [M+H] +.

1H NMR (400 MHz, DMSO-d6) δ (ppm) = 8.24 (d, J = 2.0 Hz, 1H), 7.95 (dd, J = 2.0, 8.4 Hz, 1H), 7.64 (d, J = 8.4 Hz, 1H), 3.65 (d, J = 4.8 Hz, 3H), 3.49 (dd, J = 7.8, 14.0 Hz, 1H), 3.42 - 3.31 (m, 1H)

Step 4:

To a mixture of 2-amino-3-(4-bromo-2-nitrophenyl)propanoic acid (3.50 g, 12.1 mmol, 1.00 eq) in THF (100 mL) and MeOH (100 mL) at 0 °C was added SnCl_2_ (11.4 g, 60.5 mmol, 1.57 mL, 5.00 eq) and NaOAc (9.93 g, 121 mmol, 10.0 eq). The mixture was allowed to stir, gradually warming to 20 °C for 1.5h. Et_3_N (12.2 g, 121 mmol, 16.8 mL, 10.0 eq) and Boc_2_O (7.93 g, 36.3 mmol, 8.34 mL, 3.00 eq) were then added and the mixture was stirred at 20 °C for 12h. The mixture was then filtered and the filtrate was concentrated under reduced pressure. To the residue was added water (100 mL) and then extracted with EtOAc (2 x 100 mL). The combined organic layers were dried over Na_2_SO_4_ and concentrated under reduced pressure. The residue was first purified by flash silica gel chromatography (ISCO®; 40 g Se pa Flash® Silica Flash Column, Eluent of 0 - 20% EtOAc/Petroleum ether gradient @ 100 mL/min), then purified by SFC separation (column: DAICEL CHIRALPAK AD (250mm*50mm, 10um); mobile phase: [0.1% NH_4_OH IPA]; B%: 25% - 25%, min). (R)-tert-butyl (7-bromo-1-((tert-butoxycarbonyl)oxy)-2-oxo-1,2,3,4-tetrahydroquinolin-3-yl)carbamate (100 mg) and (S)-tert-butyl (7-bromo-1-((tert-butoxycarbonyl)oxy)-2-oxo-1,2,3,4-tetrahydroquinolin-3-yl)carbamate (1000 mg) were obtained as white solids. LCMS: tr = 0.613 min, m/z = 300 [M-156] +.

SFC peak 1 (R-isomer), Rt = 0.810 min, *ee* = 97.9%.

SFC peak 2 (S-isomer), Rt = 0.996 min, *ee* = 99.5%.

1H NMR (400 MHz, MeOD-d4) δ (ppm) = 7.30 - 7.14 (m, 3H), 4.59 - 4.45 (m, 1H), 3.15 - 3.02 (m, 2H), 1.56 (s, 9H), 1.47 (s, 9H)

Step 5:

To a solution of (R)-tert-butyl (7-bromo-1-((tert-butoxycarbonyl)oxy)-2-oxo-1,2,3,4-tetrahydroquinolin-3-yl)carbamate (two batches reunited, 270 mg, 590 μmol, 1.00 eq) in EtOAc (0.5 mL) was added HCl/EtOAc (3 mL), then the mixture was stirred at 25 °C for 12h. The mixture was concentrated under reduced pressure, and the residue was purified by prep-HPLC (basic condition, column: Phenomenex C18 75*30mm*3um; mobile phase: [water (NH_4_OH+NH_4_HCO_3_ pH: 10-11)-ACN]; B%: 2% - 42%, 10 min). Compound BV3572-R (110 mg, 419 μmol, 71.0% yield, 98.3% purity) was obtained as a white solid. LCMS: tr = 0.370 min, m/z = 256 [M+1] +.

HPLC: RT = 1.78 min, purity = 98.3%.

SFC: RT = 1.54 min, ee = 97.2%.

1H NMR (400 MHz, DMSO-d6) δ (ppm) = 7.29 (s, 1H), 7.21 - 7.14 (m, 2H), 3.52 (dd, J = 6.0, 12.8 Hz, 1H), 2.99 (dd, J = 6.0, 15.6 Hz, 1H), 2.69 (dd, J = 12, 15.4 Hz, 1H)

### *Mtb* strains

All strains were cultured from frozen stocks in Middlebrook 7H9 medium (BD Difco) supplemented with glycerol 0.2% (Euromedex), Tween 80 0.05% (Sigma-Aldrich) and oleic acid-albumin-dextrose-catalase 10% (OADC, BD Difco) except if otherwise indicated.

Recombinant *Mtb* strains expressing an enhanced green fluorescent protein (GFP) were obtained by transformation with an integrative plasmid containing the *Aequoria Victoria egfp* gene(43).

H37Rv Δ*whiB7* was generated using the ORBIT recombineering system, as previously described (44). Briefly, H37Rv harboring the plasmid pKM444 (Addgene #108319) was co-electroporated with 1 μg of the attP-containing oligonucleotide targeting *whiB7* gene (whiB7-<u>attP</u>: TGAGCCGGCTCGCGCCGGCGGGCGCACCATCGCGGG<u>GGTTTGTACCGTACACCACTGAGACCGCGG</u> <u>TGGTTGACCAGACAAACC</u>GTCCTGTTTCACCTGCTTCCTGGTCTGGTGGCGGTT) and 200 ng of the attB-containing plasmid pKM464 (payload plasmid, Addgene #108322). Transformants were selected on 7H11 agar plates containing hygromycin 50 µg/mL and deletions were confirmed by PCR and Sanger sequencing of the 5′ and 3′ ends of the plasmid insertion with the primer couples (i) MTB-WhiB7-FC (5’-GTTCGCGAGATCGACGAGATGCTG-3’) and pKM464-FC (5’-cctggcagtcgatcgtacgc-3’) and (ii) MTB WhiB7-RC (5’-ATTGACAACAACGCATTTGGTGTC-3’) and pKM464-RV (5’-cagatctggtaccgctagcgg-3’).

*Mtb* H37Rv constitutively overexpressing HisC, AlaA, or Alr was obtained by transformation with plasmids pMV261-*hisC*, -*alaA*, or -*alr* and cultivated in the presence of kanamycin 25 µg/mL. As a control, H37Rv was also transformed with the empty pMV261 vector. Plasmids pMV261-*hisC*, *-alaA*, and -*alr* were constructed by NEBuilder HiFi DNA assembly method (New England Biolabs). Briefly, pMV261 was amplified by PCR in 2 segments using primer couples (i) pMV-F1 (5’-GAAGACAATTGCGGATCCAGCTGC-3’) and pMV-R1 (5’-CGCGGGTACGAGCCACAC-3’) and (ii) pMV-F2 (5’-GTGTGGCTCGTACCCGCG-3’) and pMV-R2 (5’-TTGCGAAGTGATTCCTCCGGATCG-3’). Genes *hisC*, *aspC*, and *alr* were each amplified from the genome of H37Rv using the following primers.

pMV-hisC-F 5’-CGATCCGGAGGAATCACTTCGCAATGACCAGGTCCGGACACCCG-3’

pMV-hisC-R 5’-GCAGCTGGATCCGCAATTGTCTTTCATGGCGCTCCTACAGGACTG-3’

pMV-aspC-F 5’-CGATCCGGAGGAATCACTTCGCAATGGCCGTGGACAACGATGG-3’

pMV-aspC-R 5’-GCAGCTGGATCCGCAATTGTCTTCTATTGCCGGTAACTGACCAGGAAG-3’

pMV-alr-F 5’-CGATCCGGAGGAATCACTTCGCAATGGCCGTGAAACGGTTCTGGGAGAA-3’

pMV-alr-R 5’-GCAGCTGGATCCGCAATTGTCTTTCAACGGTTTTCAGCCTCGC-3’

### Extracellular antibacterial activity

Stock solutions of BVL3572S and analogues were prepared in DMSO at 34 mM (10 mg/mL). Aliquots were stored at -20 °C. Compounds were dispensed into black Greiner 384-well clear bottom polystyrene plates (Greiner Bio-One 781091) using an HP D300e digital dispenser. DMSO was added to relevant wells to ensure a final concentration of 0.6% across all wells.

The assay plates were inoculated with 50 µL/well of GFP-expressing *Mtb* H37Rv or derivatives at an optical density (OD_600 nm_) of 0.02 and incubated at 37 °C for 5 days. For complementation studies, L-His, L-Ala, or a combination of both were added at a final concentration of 3 mM and PLP was supplemented at 100 µM. GFP fluorescence (λex = 485 nm/λem = 535 nm) was read using the Ensight Multimode plate reader (PerkinElmer, USA). IC50 values were calculated by fitting fluorescence values for each concentration on GraphPad Prism 9 (GraphPad Software, Inc., San Diego, CA). When indicated, MIC_90_ values correspond to the first well for which normalized fluorescence values were ≤ 10% of the maximum fluorescence of drug-free wells. All assays were conducted with two technical duplicates per biological replicate, with a total of three biological replicates.

### Intracellular antibacterial activity

THP-1 cells were transfected with Incucyte® CytoLight Red Lentivirus (EF-1 Alpha promoter, puromycin selection, Sartorius 4482) according to the protocol described in (45). Transfected THP-1 cells were then cultured at a density of 2x10^5^ to 1x10^6^ cells/mL in RPMI medium (Thermo Fisher 61870036) supplemented with 10% fetal bovine serum (FBS, Gibco 10500) at 37 °C, 5% CO_2_. A single-cell bacterial suspension of *Mtb* H37Rv pEG200 (expressing a luciferase operon) was prepared for infection (46,47). Briefly, an exponentially growing culture was centrifuged at 3,200 x g for 5 min and the pellet resuspended in RPMI-FBS before centrifugation at 100 x g to remove clumps. The supernatant was passed through a cell strainer with a porosity of 10 µm and the OD was measured. THP-1 cells (5 mL of 5x10^6^ cells) were simultaneously infected by adding the bacterial suspension at a multiplicity of infection of 10 and differentiated with 40 ng/mL phorbol 12-myristate-13-acetate (Sigma P1585). The cells were incubated for 4 h at 37 °C, 5% CO_2_ and the adherent cell monolayer was detached with a cell scraper. The cells were washed 4 times by centrifugation at 300 x g and the pellet resuspended in 25 mL of RPMI-FBS. Opaque white 384-well assay plates were prepared as described above for the extracellular assay. Plates were inoculated with 50 µL of the cell suspension (1x10^4^ cells/well) and incubated for 5 days at 37 °C, 5% CO_2_. For complementation studies, plates were supplemented with 80 µM of L-His, L-Ala, or both. The antibacterial activity of the compounds was determined by measuring luminescence of intracellular bacteria. The experiment was performed in triplicate.

### Time-kill assay

Cultures were treated with various concentrations of BVL3572S (stock solution prepared in DMSO) in non-ventilated 25 cm² tissue-culture flasks and each contained a fixed DMSO concentration (0.3%). An untreated control containing only DMSO was included. The flasks were inoculated with 10 mL of *Mtb* H37Rv cultures diluted to an OD_600 nm_ of 0.005 (theoretically corresponding to 5 x 10^5^ CFU/mL) and incubated at 37 °C without shaking.

Cultures from each flask were plated on square plates containing Middlebrook 7H10 solid medium (BD) supplemented with glycerol 0.2%, Tween 80 0.05%, and oleic acid-albumin-dextrose-catalase 10% on days 0, 2, 4, 7, 14, and 21. On day 0, only 3 cultures were plated whereas on other days, all cultures were plated. Before plating, cultures were 10-fold serially diluted (up to 10^-7^) and 10 µL of each serial dilution (including the non-diluted culture) were plated. CFUs were counted after incubating the plates for up to 3 weeks at 37 °C. The lowest possible number of countable CFUs (limit of detection) is 1 when plating 10 µL of a non-diluted culture, which amounts to a log_10_ CFU/mL of 2. Three biological replicates were performed.

### Transcriptomics

Exponentially growing *Mtb* H37Rv cultures (25 mL; OD_600 nm_ 0.7) were exposed to 20 µM of BVL3572S or 0.5% DMSO for 6 h. The treatment was performed for independent biological duplicates. RNA extraction and transcriptomics analysis were performed as previously described (48). Briefly, 10 mL of the cultures were harvested by centrifugation at 3,500 x g for 5 min, resuspended in 1 mL of RNApro^TM^ (FastRNA Pro Blue Kit, MP biomedicals) and homogenized in impact-resistant 2 mL tubes containing 0.1 mm silica spheres (Lysing Matrix B, MP biomedicals) using a FastPrep FP120 cell disrupter (Thermo Fisher Scientific) at 6.0 Hz for 40 s. The lysates were centrifuged at 12,000 x g and RNA was purified according to the manufacturer’s instructions. Ribosomal RNA (rRNA) was depleted using the QIAseq FastSelect 5S/16S/23S kit (Qiagen). Illumina sequencing libraires were prepared with the TruSeq RNA sample preparation kit version 2.0 rev. A (Illumina Inc.) with a unique index for each cDNA library. cDNA libraries were sequenced using an Illumina NextSeq 500 system (Illumina Inc.) in high-output mode. All samples were multiplexed on a single flow-cell lane and sequenced in single-read sequencing mode with read lengths of 150 bp. Raw reads were processed with Illumina quality control tools using default parameters. Sequences shorter than 50 bp, containing ambiguous bases (‘Ns’), and/or with a mean quality score lower than 30 were removed using PRINSEQ (http://prinseq.sourceforge.net/index.html). Next, rRNA-specific reads were excluded by aligning all reads on *Mtb* rRNA sequences using Bowtie2 (http://bowtie-bio.sourceforge.net/bowtie2/index.shtml). Downstream RNA-seq analysis was performed using the SPARTA open-source software package with default settings (https://sparta.readthedocs.io/en/latest/), yielding expression data for 4036 genes.

### Selection of spontaneous resistant mutants

Plates containing Middlebrook 7H10 or 7H9 solid medium supplemented with glycerol 0.2%, Tween 80 0.05% and oleic acid-albumin-dextrose-catalase 10% were prepared with BVL3572S alone (7H9) or in the presence of L-His 3 mM (7H9) or L-Ala 3 mM (7H10). 7H9 solid medium was used for mutant selection with L-His or alone because we suspected our 7H10 stock to contain trace amounts of L-Ala. Plates were inoculated with 10^6^, 10^7^, or 10^8^ CFUs of a weeks-old culture of *Mtb* H37Rv (OD_600 nm_ > 1) and incubated at 37 °C for at least 4 weeks. Individual colonies were picked, cultured in the presence of the corresponding amino acid if applicable, and the growth inhibitory activity of BVL3572S was determined as indicated above. Stocks of each culture were prepared and stored at -80 °C.

For genomic DNA sequencing, 7 mL of exponentially growing cultures were harvested by centrifugation at 3,500 x g for 5 min, resuspended in 500 µL of UltraPure™ Phenol:Chloroform:Isoamyl Alcohol (Invitrogen), and homogenized in impact-resistant 2 mL tubes containing 0.1 mm silica spheres (Lysing Matrix B, MP biomedicals) using a FastPrep FP120 cell disrupter (Thermo Fisher Scientific) at 6.0 Hz for 40 s. Samples were mixed with 500 µL TE buffer (10 mM Tris pH 8.5, 1 mM EDTA) and incubated for 5 min at room temperature before centrifugation (10 min, 10,000 rpm, 4 °C). The aqueous phase (400 µL) was transferred to a new tube, to which 80 µL sodium acetate (4 M, pH 5.5) and 800 µL of 100% cold ethanol were added. DNA was precipitated overnight at - 20 °C. After centrifugation (30 min, 12,000 rpm, 4 °C), the supernatant was removed, and the pellet was washed twice with 700 µL of 70% cold ethanol followed by centrifugation (15 min, 12,000 rpm, 4 °C). Pellets were air-dried for around 30 min and resuspended in 35 µL nuclease-free water. DNA was quantified by Qubit dsDNA BR assay kit (ThermoFisher Scientific) and sent to Novogene for library preparation and whole genome sequencing by Illumina paired-end joining.

### Construction of *E. coli* BW25113 Δ*hisC* (pBAD18Ω*hisC*_Mtb_)

Deletion of *hisC* in *E. coli* BW25113 was obtained by phage P1 transduction of the Km^R^ cassette of the *hisC* deletion mutant from the Keio collection (49,50). Kanamycin 50 µg/mL was used for selection of transductants. BW25113 Δ*hisC*::Km^R^ clones were transformed with the FLP recombinase expression plasmid pFLP2 to remove the Km^R^ cassette. Transformants were selected with ampicillin 200 µg/mL at 37 °C. Approximately 30 independent colonies were patched in parallel onto LB agar containing either kanamycin or ampicillin and incubated at 37°C to identify recombinants that had undergone FLP-mediated excision of the Km^R^ cassette. Clones that exhibited a Km-sensitive, Amp-resistant phenotype were subsequently streaked on LB agar supplemented with 5% sucrose at 37 °C to eliminate pFLP2. Resulting sucrose-resistant colonies were patched in parallel onto LB agar with and without ampicillin to identify Amp-sensitive clones, corresponding to the cured BW25113 Δ*hisC* strain. Several Amp^S^ clones were isolated on LB medium, and loss of both the *hisC* gene and Km^R^ cassette was confirmed by PCR using primers located approximately 200 bp upstream and downstream of the *hisC* locus. Verified deletion strains were stored as glycerol stocks.

To construct pBAD18Ω*hisC*_Mtb_, the *hisC_Mtb_* gene construct was digested from pET28aΩ*hisC*_Mtb_ (see below) using XbaI and HindIII and cloned into plasmid pBAD18 between the XbaI and HindIII restriction sites. The plasmid or the empty vector was transformed into BW25113 WT and BW25113 Δ*hisC*. Transformants were selected with ampicillin 200 µg/mL.

### Plating efficiency assay for *E. coli*

*E. coli* BW25113 and derivatives were streaked from frozen stocks on solid LB medium (containing ampicillin 100 µg/mL if pBAD18 was present) and incubated overnight at 37 °C. Liquid cultures in M9 medium, containing ampicillin 100 µg/mL if needed and L-His 80 µM to allow growth of auxotrophs, were inoculated with a single isolated colony and incubated overnight at 37 °C under agitation. The OD_600 nm_ of all cultures was adjusted to 1.0 and 10-fold serial dilutions (up to 10^-6^) were prepared in M9 medium. The suspensions (5 µl) were plated on M9-agar plates in the absence or presence of L-His 80 µM or L-arabinose 0.1%. Plates were incubated for 24 h at 37 °C before being imaged. Pictures shown are representative of at least 2 biological replicates.

### MIC determination and checkerboard assay for *E. coli*

*E. coli* BW25113 and derivatives were streaked from frozen stocks on solid LB medium (containing ampicillin 100 µg/mL if pBAD18 was present) and incubated overnight at 37°C. Liquid cultures in M9 medium, containing ampicillin 100 µg/mL if needed and L-His 80 µM to allow growth of auxotrophs, were inoculated with a single isolated colony and incubated overnight at 37 °C under agitation.

For MIC determination, assay plates were prepared as described for *Mtb* H37Rv except that U-bottom 96-well plates were used. Plates were inoculated with an overnight *E. coli* BW25113 culture in M9 medium at an OD_600 nm_ of 0.002. L-His, L-Ala, or a combination of both were supplemented at 80 µM each. MIC was determined as the lowest concentration of drug that led to no visible growth after 24 h of incubation at 37 °C.

For checkerboard assays, flat-bottom 96-well plates were distributed with 2-fold serial dilutions of BVL3572S or rifampicin (stock solutions prepared in DMSO) using an HP D300e digital dispenser. L-arabinose (stock solution prepared in water) was serially diluted and distributed manually. Plates were inoculated with overnight *E. coli* cultures in M9 medium at an OD_600 nm_ of 0.002 and incubated for 24 h at 37 °C. OD_600 nm_ was read using a Tecan M Plex Infinite 200 PRO plate reader. When required, L-His was supplemented at 80 µM each. Absorbance values for each concentration were fitted on GraphPad Prism 9 (GraphPad Software, Inc., San Diego, CA).

### Enzyme inhibition assays

The enzymatic assay of HisC was performed in the reverse direction as previously described (51). Enzymatic activity was measured using 0.125 µM (5 µg/mL) HisC in 0.13 M triethanolamine (pH 8.5) buffer containing 3.4 mM 2-oxoglutarate; the reaction was started by adding 1.7 mM histidinol phosphate. Upon incubating this 100 µL reaction mixture for 90 min at 37 °C to allow formation of L-glutamate and imidazole acetol phosphate, the reaction was stopped by adding 900 µL of 1.43 M NaOH and incubating for 20 min at 45 °C. Enolized imidazole acetol phosphate was spectrophotometrically detected at 280 nm. To measure the inhibitory activity of BVL3572S, the compound was included in the reaction at various concentrations prior to the addition of HisC.

### Purification of *Mtb* HisC

The *hisC*_Mtb_ gene was amplified from genomic DNA using Phusion high fidelity polymerase (New England Biolabs) with primers 5’-GTGCGGCCGCAAGCTTTGGCGCTCCTACAGGACTG-3’ and 5’-AGGAGATATACCATGGTTATGACCAGGTCCGGACACC-3’. PCR fragments were cloned into plasmid pET28a between the HindIII and NcoI restriction sites using the In-Fusion Cloning (Takara Bio) method. The pET28aΩ*hisC*_Mtb_ plasmid was cloned into *E. coli* BW25113 for protein production.

LB medium containing kanamycin 25 µg/mL was inoculated with an overnight preculture of *E. coli* BL21(DE3) pET28aΩ*hisC*_Mtb_ and grown at 37 °C under agitation up to an OD_600 nm_ of 0.8–1. Protein expression was induced with 0.5 mM IPTG for 3 h at 37 °C under agitation. Cells were harvested by centrifugation (4,500 x g, 30 min, 4 °C), and the pellet was resuspended in lysis buffer (20 mM Tris pH 7.5, 150 mM NaCl, 20 mM imidazole) supplemented with protease inhibitors, DNase I, and SDS (final 0.1%). Bacterial lysis was performed by sonication on ice. Cell debris was removed by centrifugation (10,000 x g, 30 min, 4 °C) and the clarified lysate was loaded onto a pre-equilibrated HisTrap™ HP 5 mL column (Cytivia) at 2 mL/min. The column was washed with buffer (20 mM Tris pH 7.5, 150 mM NaCl, 20 mM imidazole) and proteins were eluted with 250 mM imidazole in the same buffer. Eluted fractions were pooled and dialyzed overnight at 4 °C against the lysis buffer. The dialyzed sample was concentrated using Millipore Amicon Ultra-4 centrifugal filters (30 kDa cutoff), and supplemented with 10% glycerol except when samples were used for protein crystallization.

### Structure determination of the HisC-BVL3572S complex

Crystals of HisC_Mtb_ bound to PLP and MES were obtained using the hanging-drop vapor diffusion method at 298 K with a protein concentration of 7.5 mg/mL. The crystallization solution contained 100 mM ammonium sulfate, 10 mM Tris (pH 7.0), 75 mM NaCl, 15% (w/v) PEG MME 5000, 50 mM MES, and 25 µM PLP. The HisC-BVL2572S complex was generated by soaking crystals in the same solution supplemented with 0.5 mM BVL3572S, while maintaining the final DMSO concentration below 1% (v/v). Soaking times ranging from 1 to 40 s were tested. Notably, the presence of PLP in the soaking solution was essential to trap the compound in the protein active site. Crystals were cryoprotected with 20% (v/v) glycerol and flash-cooled in liquid nitrogen for X-ray data collection at the Proxima 2A beamline (SOLEIL synchrotron).

Diffraction data were processed (scaling and indexing) using the XDSME pipeline. The structure was solved by molecular replacement with MOLREP (52), using the HisC-PLP structure (PDB code: 4R8D) as a search model. Refinement was carried out through iterative cycles of REFMAC5 (53) and manual rebuilding in Coot (54). Data collection and refinement statistics are provided in Supplementary Table 2.

An omit electron density map was calculated by excluding BVL3572S from the model, randomly perturbing the remaining atom coordinates (0-0.3 Å), recalculating structure factors, and computing the residual map. Ligand-protein interactions in the HisC*_Mtb_*-BLV3572S complex were analyzed using the LigPlot+ software (55).

### Detection of PLP-BVL3572S adduct

Purified HisC_Mtb_ (21 µM) was mixed with BVL3572S (21 µM) and PLP (50 µM) for 60 min at 25°C. The mixture was loaded onto a HisTrap™ column and the flow-through was harvested. Controls containing PLP alone, BVL3572S alone, or a mix of both were prepared at the indicated concentrations. LC–MS analysis was performed on an Alliance e2695 system with a 2998 PDA detector (Waters) coupled to an Altus SQ mass detector (PerkinElmer). Electrospray ionization was applied in positive mode under the following settings: cone voltage 49 V, capillary voltage 3.1 kV, source temperature 150 °C, and desolvation temperature 600 °C. Separation was carried out on a C18 Alltima column (250 × 4.6 mm, 10 μm; Grace) at 0.5 mL min⁻¹, using a gradient from H₂O/CH₃OH (9:1, 0.1% formic acid) to H₂O/CH₃OH (1:9, 0.1% formic acid). Data were acquired and processed with Empower 3 software. Calibration was performed by first injecting PLP alone, BVL3572S alone, and a mix of both.

### LC-MS analysis of BVL3572S-induced remodeling of amino acid ^13^C-labeling

*Mtb* H37Rv was cultured up to an OD_600 nm_ of 0.3 in Sauton medium containing NH_4_Cl (instead of aspartate) as the nitrogen source supplemented with glycerol 0.2%, Tween 80 0.05% and oleic acid-albumin-dextrose-catalase 10%. The culture was treated with BVL3572S (1 µM) or DMSO for 2 h at 37 °C before addition of ^13^C-glycerol 0.2% and further incubated for 16 h. The OD_600 nm_ of the cultures was between 0.41 to 0.43 (in the presence of treatment) and 0.46 (without treatment). The cultures were centrifuged (3,500 x g, 5 min) and the pellet resuspended in HCl 6 N before a 10 h incubation at 100 °C. Samples were then lyophilized.

Analyses were performed on an LC-MS platform consisting of a Thermo Scientific™ Vanquish™ Focused UHPLC Plus system coupled to a Thermo Scientific™ Orbitrap Exploris™ 120 hybrid quadrupole-Orbitrap™ mass spectrometer equipped with a heated electrospray ionization (HESI) probe.

Chromatographic separation of amino acids was carried out on a SUPELCO PFP column (150 × 2.1 mm, 5 µm) with a HS F5 guard column (20 × 2.1 mm, 5 µm). The column was maintained at 30 °C, and the flow rate was set at 0.25 mL/min. The solvent system consisted of (A) 0.1% formic acid in water and (B) 0.1% formic acid in acetonitrile. The gradient was adapted from the method of Boudah et al. (56) and was as follows: 0 min, 2% B; 2 min, 2% B; 10 min, 5% B; 16 min, 35% B; 20 min, 100% B; 24 min, 100% B. The column was then equilibrated for 6 min at the initial conditions before the next sample was analyzed. The injection volume was 5 µL, and the autosampler temperature was maintained at 4 °C.

Mass detection was performed in positive electrospray ionization (ESI+) mode. The mass spectrometer settings were as follows: spray voltage, 3.5 kV; capillary temperature, 325 °C; desolvation temperature, 350 °C; and maximum injection time set to Auto. Nitrogen was used as sheath gas (50 arbitrary units) and as auxiliary gas (10 arbitrary units). The automatic gain control (AGC) target was set to 1 × 10^6^, and a resolution of 60,000 was used over the m/z 50– 750 range. MS analyses were performed in Full Scan mode. Data acquisition was carried out using Thermo Scientific Xcalibur software. Metabolites were identified by extracting their exact masses with a tolerance of 10 ppm. Data processing was performed with Skyline software.

Values were normalized to untreated controls to compute percent change in ^13^C labeling under drug treatment and further adjusted using the optical density difference between treated and untreated cultures.

### Tn-seq analysis

A pooled transposon mutant library of *Mtb* H37Rv, constructed by transduction with the MycoMar T7 phage (57,58) and containing more than 20,000 mutants, was pre-cultured in 5 mL of 7H9 medium supplemented with Tween 80 0.05%, albumin-dextrose-catalase 10% (ADC, BD Difco) and kanamycin (40 µg/mL) for 5 days at 37 °C. The culture was centrifuged (3,000 rpm, 10 min) and the pellet was washed with 15 mL of Dulbecco’s phosphate buffered saline (DPBS 1X, Gibco) with Tween 80 0.05%. Bacteria were resuspended in 25 mL of fresh medium without kanamycin and cultured for 2 days at 37 °C, until an OD_600 nm_ of approximately 0.4 was reached. The culture was used to inoculate 10 flasks at an OD_600 nm_ of 0.01 in 25 mL of medium. The flasks were incubated for 3 days at 37 °C until an OD_600 nm_ of 0.2 was reached. Flasks were separated into 2 groups of 5 and each group was treated with either BVL3572S 27 µM or DMSO 0.8%. After 15 days at 37 °C, 1 mL of each culture was centrifuged (10,000 rpm, 2 min), washed with DPBS 1X supplemented with Tween 80 0.05% and diluted in fresh medium in order to obtain approximately 5 x 10^5^ bacteria/mL in 25 mL. These cultures were incubated for 7 days at 37 °C up to an OD_600 nm_ of approximately 0.6, then 12 mL of each culture were centrifuged (3000 rpm, 10 min), and the pellets were conserved at -20 °C until DNA extraction.

Genomic DNA was extracted (Quick-DNA^TM^ Fungal/Bacterial Miniprep Kit, Zymo Research), and transposon-genome junctions were amplified by PCR as previously described (58). An index specific to each sample was added during this PCR, allowing all samples to be pooled to create the Tn-seq library (TruSeq^MC^ DNA Nano). This library was sequenced by Illumina paired-end sequencing to determine the transposon mutant composition of each sample. Resulting reads were mapped onto the *Mtb* H37Rv reference genome. For each sample, a table was generated containing the number of read for each TA insertion site (58). Data were processed using the TRANSIT pipeline (59). Insertion counts at each TA site were normalized using timed total reads (TTR) normalization. The resampling test module in TRANSIT was used to identify genes conditionally affecting fitness in the presence of BVL3572S treatment. Significant differences in read counts between BVL3572S-free and BVL3572S-treated conditions were assessed by comparison to a resampling distribution generated through random permutation of TA site counts within each genetic locus across all datasets. P values were calculated as the proportion of values from 10⁶ permutations that were more extreme than the observed result. To account for multiple comparisons, P values were adjusted using the Benjamini–Hochberg procedure to obtain false discovery rates. TnSeq fold changes were expressed as log₂ ratios of normalized read counts between the BVL3572S-treated and BVL3572S-free libraries.

### CRISPRi screen

CRISPRi screens were performed and analyzed as previously described with slight modifications (3,60). Briefly, 3 x 1mL *Mtb* CRISPRi library aliquots (RLC12, Addgene 163954) were thawed and inoculated in 10 mL of 7H9 containing kanamycin 20 µg/mL (OD_600 nm_ = 0.1). After a 5-day incubation at 37 °C, cultures were centrifuged to get rid of clumps (100 x g, 2 min) and the supernatant was diluted to an OD_600 nm_ of 0.1 in 40 mL of 7H9 containing kanamycin and anhydrotetracycline (ATc) at 100 ng/mL. Target pre-depletion was performed for 5 days after which, medium was renewed by centrifuging the cultures (3,500 x g, 5 min) and resuspending the pellet in 7H9 before dilution to an OD_600 nm_ of 0.05 in 20 mL of kanamycin- and ATc-containing 7H9. Treatment was initiated by adding sub-MIC BVL3572S concentrations alone (0.85, 0.43, or 0.21 µM), with L-His 3 mM (0.43, 0.21, or 0.11 µM), with L-Ala 3 mM (1.70, 0.85, or 0.43 µM), or with 3 mM each of both L-His and L-Ala (13.63, 6.81, or 3.41 µM) to triplicate cultures. Antibiotic-free controls were prepared by adding DMSO 0.1% to cultures containing the corresponding amino acid. Cultures were outgrown for 14 days. On days 5 and 10 of outgrowth, medium was renewed by centrifuging the cultures and resuspending the pellets in medium before dilution to an OD_600 nm_ of 0.1 in 7H9 containing kanamycin, ATc, and if applicable, BVL3572S and amino acid at the indicated concentrations. On day 14 of outgrowth, the impact of BVL3572S treatment on each culture was determined by measuring OD_600 nm_ (the OD_600 nm_ of untreated cultures was approximately 1). Cultures with partially drug inhibitory BVL3572S concentrations (growth reduction of 20 to 50%) and the corresponding controls were processed for genomic DNA extraction. Retained BVL3572S concentrations were 0.43 µM for the drug alone, 0.21 µM in the presence of L-His, 1.7 µM in the presence of L-Ala, and 6.81 µM in the presence of both amino acids. For BVL3572S + L-Ala, only 2 samples were successfully processed, one of which was subsequently sequenced twice to generate 3 replicates.

Genomic DNA was isolated from 20 mL of selected cultures using the CTAB-lysozyme method as previously described (60) and quantified using the Qubit dsDNA BR assay kit (ThermoFisher Scientific). The sgRNA-encoding region was amplified from genomic DNA exactly as previously described (60). Amplicons (∼230 bp) were purified using NucleoMag magnetic beads (Macherey-Nagel 744970) using a double-sided size selection (bead:sample ratio of 0.6 followed by 1.0) and quantified using the Qubit dsDNA BR assay kit (ThermoFisher Scientific). Amplicon size and purity were visualized on a 1 % agarose gel and an Agilent 2100 bioanalyzer. For a more precise quantification, a qPCR assay was performed using the KAPA SYBR FAST qPCR kit (Roche KK4600). Individual PCR amplicons were multiplexed into 4 nM pools and sequenced on an Illumina Novaseq PE150 platform.

sgRNA sequences were extracted from single end reads using *cutadapt* (61) between sequences [GTGATAGATATAATCTGGGA…GTTTTTGTAGCTCGAAAGAAG]. Extracted sequences shorter than 4 nucleotides were discarded after trimming. sgRNA sequences were mapped and quantified using MAGeCK (62) *count* function. MAGeCK *test* function was used to determine sgRNA and gene level enrichment or depletion, using alphamedian L2FC method and Negative (control) sgRNA list for normalization (60).

### Checkerboard assay

All stock solutions were prepared at 10 mg/mL in DMSO 100% except D-cycloserine, which was solubilized in water. Checkerboard assays were performed exactly as described above for the extracellular antibacterial activity assay except that 2-fold serial dilutions of BVL3572S and companion drugs were distributed vertically and horizontally, respectively, in 384-well plates using an HP D300e digital dispenser. When required, clavulanic acid was distributed at a fixed concentration of 4 µg/mL using the dispenser. D-cycloserine was serially diluted and distributed manually. Because of its instability, meropenem was supplemented to the test plates at the indicated concentrations on day 3. The assay plates were inoculated with 50 µL/well of GFP-expressing *Mtb* H37Rv at an OD_600 nm_ of 0.02 and incubated at 37 °C for 5 days. GFP fluorescence (λex = 485 nm/λem = 535 nm) was read using the Ensight Multimode plate reader (PerkinElmer, USA). When indicated, MIC_90_ values correspond to the first well for which normalized fluorescence values were ≤ 10% of the maximum fluorescence of drug-free wells. All assays were conducted with two technical duplicates per biological replicate, with a total of three biological replicates.

### Accession numbers

The electron density map, and the coordinates of the refined crystal structure were deposited to the Protein Data Bank under PDB ID: pdb_00009T0W.

## Acknowledgment

This research was supported by Inserm, CNRS, the University of Lille and the French government’s Investissements d’Avenir program (grant reference: Mustart ANR-20-PAMR-0005). We acknowledge support by the MetaboHUB infrastructure funded by the Agence Nationale de la Recherche under the France 2030 program (MetaboHUB ANR-11-INBS-0010; MetEx+ ANR-21-ESRE-0035; MetaboHUB (JVCE) ANR-24-INBS-0012). The SOLEIL synchrotron is acknowledged for access to the synchrotron-radiation facility (BAG20210875). A special thanks to Dr. Abdalkarim Tanina for his assistance and help with HisC crystallization. RW is a Research Associate at the Belgian Funds for Scientific Research (FNRS).

