## Supplementary Information for "Exploiting Vitamin B6 Dependency: BVL3572S Inhibits HisC and AlaA to Kill *Mycobacterium tuberculosis*"

**Supplementary Table 1. Spontaneous BVL3572S-resistant mutants and their genomic mutations.** Mutations in *hisC* and *alaA* are indicated in bold. Mutations present in multiple mutants are color-coded consistently.

| Mutant | Mutant selection: Amino acid | Mutant selection: BVL3572S (µM) | Genome position | Gene | Mutation | Amino acid substitution |
|---|---|---|---|---|---|---|
| **25_1c** | None | 8.5 | **402387** | ***alaA*** | **776C>A** | **Thr259Asn** |
| | | | 1129072 | *rpfB* | 982G>A | Ala328Thr |
| | | | 1466686 | *atpD* | 846C>T | Synonymous |
| | | | 2257340^2257341 | *Rv2008c* | 602dupG | Leu202fs |
| | | | 2949779 | *TB31.7* | 187T>C | Trp63Arg |
| **125H1c** | L-His | 4.3 | 400208 | *Rv0336* | 17C>T | Ala6Val |
| | | | **402355** | ***alaA*** | **808G>A** | **Ala270Thr** |
| | | | 1457326 | *hemK* | 762T>C | Synonymous |
| **5H1c** | L-His | 17 | **402355** | ***alaA*** | **808G>A** | **Ala270Thr** |
| | | | 642038 | *fadD8* | 774C>T | Synonymous |
| | | | 994059 | *Rv0892* | 207T>C | Synonymous |
| | | | 3346450 | *Rv2989* | 304C>T | Arg102Cys |
| | | | 3903190 | *Rv3484* | 113T>C | Leu38Pro |
| **5H3c** | L-His | 17 | 400208 | *Rv0336* | 17C>T | Ala6Val |
| | | | **402355** | ***alaA*** | **808G>A** | **Ala270Thr** |
| | | | 1457326 | *hemK* | 762T>C | Synonymous |
| **25A6b** | L-Ala | 8.5 | 400208 | *Rv0336* | 17C>T | Ala6Val |
| | | | 1457326 | *hemK* | 762T>C | Synonymous |
| | | | **1801335** | ***hisC*** | **440C>T** | **Ala147Val** |
| | | | 2601651 | *PE23* | 921G>A | Synonymous |
| **5A1** | L-Ala | 17 | 169656 | *Rv0143c* | 527C>T | Ala176Val |
| | | | 642038 | *fadD8* | 774C>T | Synonymous |
| | | | 994059 | *Rv0892* | 207T>C | Synonymous |
| | | | **1801242** | ***hisC*** | **347C>T** | **Pro116Leu** |
| | | | 3258369 | *ppsC* | 2691dupC | Glu898fs |
| | | | 3346450 | *Rv2989* | 304C>T | Arg102Cys |
| | | | 3903190 | *Rv3484* | 113T>C | Leu38Pro |
| **5A2** | L-Ala | 17 | 642038 | *fadD8* | 774C>T | Synonymous |
| | | | 994059 | *Rv0892* | 207T>C | Synonymous |
| | | | 1273988 | *Rv1146* | 634G>A | Val212Ile |
| | | | **1801318** | ***hisC*** | **423G>T** | **Trp141Cys** |
| | | | 3203254 | *Rv2893* | 835A>G | Thr279Ala |
| | | | 3346450 | *Rv2989* | 304C>T | Arg102Cys |
| | | | 3903190 | *Rv3484* | 113T>C | Leu38Pro |
| **10A1** | L-Ala | 34 | 642038 | *fadD8* | 774C>T | Synonymous |
| | | | 994059 | *Rv0892* | 207T>C | Synonymous |
| | | | **1801254** | ***hisC*** | **359C>T** | **Ala120Val** |
| | | | 2062847 | *bacA* | 1882C>A | Synonymous |
| | | | 3258369 | *ppsC* | 2691dupC | Glu898fs |
| | | | 3346450 | *Rv2989* | 304C>T | Arg102Cys |
| | | | 3903190 | *Rv3484* | 113T>C | Leu38Pro |

**Supplementary Table 2.** Data collection and refinement statistics of the structure of the PLP-BVL3572S/HisC complex.

| | |
|---|---|
| **Data Collection[a]** | |
| PDB id | pdb_00009T0W |
| Synchrotron beamline | PROXIMA-2A |
| Wavelength (Å) | 0.98011 |
| Space group | $P2_12_12_1$ |
| Unit cell parameters (Å) | 67.60;101.59;115.53 |
| Range of resolution limit (Å) | 58.348-1.737 (1.767-1.737) |
| Completeness (%) | 99.6 (99.7) |
| No. of reflections | 586944 (24292) |
| No. of unique reflections | 82707 (4080) |
| Multiplicity | 7.1 (6.0) |
| $R_{meas}$[a] | 0.181 (2.682) |
| $<I/\sigma(I)>$[b] | 7.5 (0.8) |
| $CC_{1/2}$[c] | 0.995 (0.343) |
| **Refinement statistics** | |
| Number of reflections[d] | 82395 (4098) |
| Number of refined atoms | |
| Protein | 5502 |
| Water | 234 |
| Other | 95 |
| Final $R_{work}$[e]/$R_{free}$[f] (%) | 19.4/23.1 |
| RMSD bond lengths (Å) | 0.014 |
| RMSD bond angles (°) | 2.199 |
| Overall mean B factor ($Å^2$) | 31.47 |
| Compound mean B factor ($Å^2$) | 36.5 |
| Mean B factor for interacting atoms ($Å^2$) | 22.775 |
| **MolProbity statistics** | |
| Ramachandran favoured (%) | 96.27 |
| Ramachandran outliers (%) | 0.28 |
| Clash score all-atom | 2.51 (99th percentile) |
| MolProbity score | 1.43 (94th percentile) |
| **PLP-BVL3572S density fit analysis[h]** | |
| RSR | 0.077 |
| RSCC | 0.959 |
| RSZO | 4 |
| RSZD | 5.8 |

[a] $R_{meas} = \sum_{hkl} [N/(N-1)]^{1/2} \sum_i |I_i(hkl) - <I(hkl)>|/\sum_{hkl} \sum_i I_i(hkl)$ , is the multiplicity ($N$) independent $R_{merge}$, where $I_i$ is the intensity of the $i^{th}$ observation and $<I>$ is the mean intensity of the reflections.

[b] $<I/\sigma(I)>$ = mean of $I/\sigma(I)$ of unique reflections.

[c] The mean intensity correlation coefficient of half-datasets.

[d] Numbers in parentheses are the number of reflections in the free set.

[e] $R_{work} = \sum||F_{obs}| - |F_{calc}||/\sum|F_{obs}|$, where $F_{calc}$ and $F_{obs}$ are the calculated and observed structure factor amplitude, respectively.

[f] $R_{free} = \sum||F_{obs}| - |F_{calc}||/\sum|F_{obs}|$, where all reflections belong to a test set of 5% randomly selected data.

[g] RSR, RSCC, RSZO and RSZD are the real-space R factor, the real-space correlation coefficient, the Real-space $Z^{obs}$ metric and the real-space $Z^{diff}$ metric, respectively. These values were computed using EDSTATS (1).

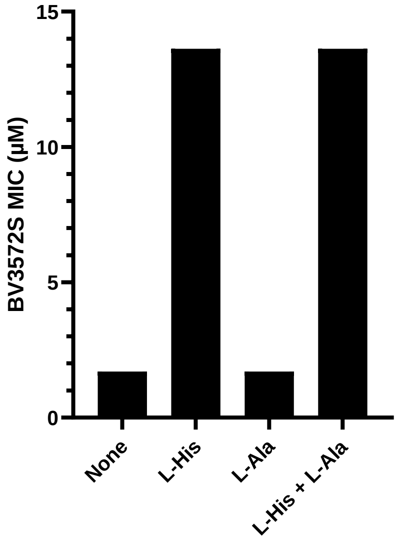


**Supplementary Fig. 1. BVL3572S targets HisC in *E. coli*. (A)** MIC determination of BVL3572S against WT *E. coli* BW25113 supplemented with indicated amino acids at 80 μM. Data represent three biological replicates.

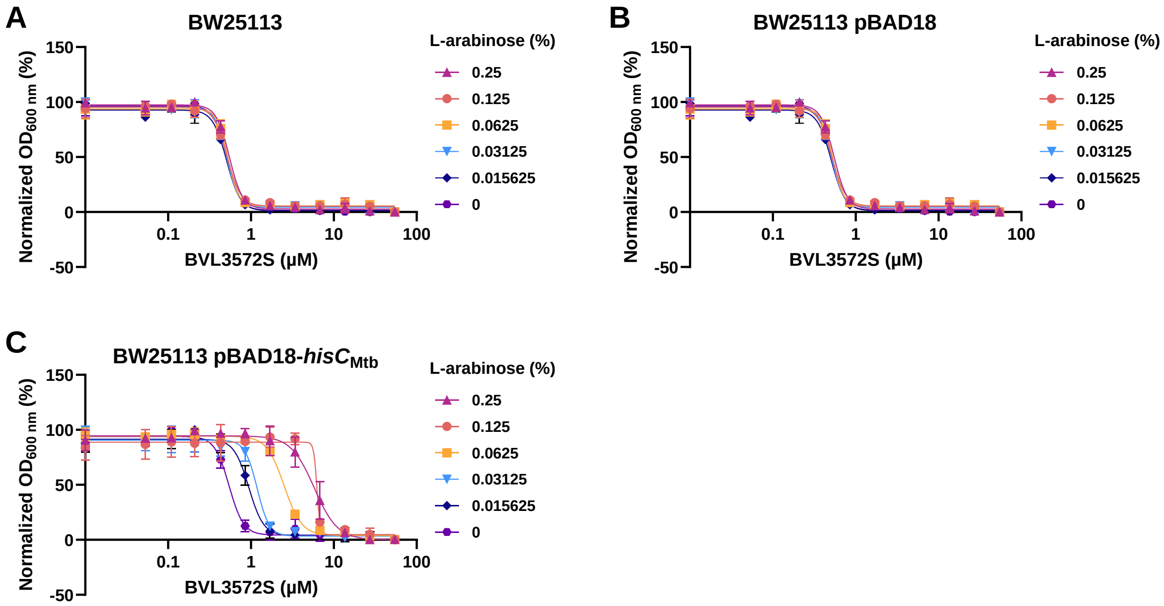


**Supplementary Fig. 2. Effect of L-arabinose on BVL3572S growth inhibition in *E. coli*. (A–C)** Growth inhibition by BVL3572S across increasing L-arabinose concentrations in **(A)** WT BW25113, **(B)** BW25113 (pBAD18) (empty vector), and **(C)** BW25113 (pBAD18Ω*hisC*$_{\text{Mtb}}$). Data shown are the mean ± SD of two technical replicates and are representative of three biological replicates.

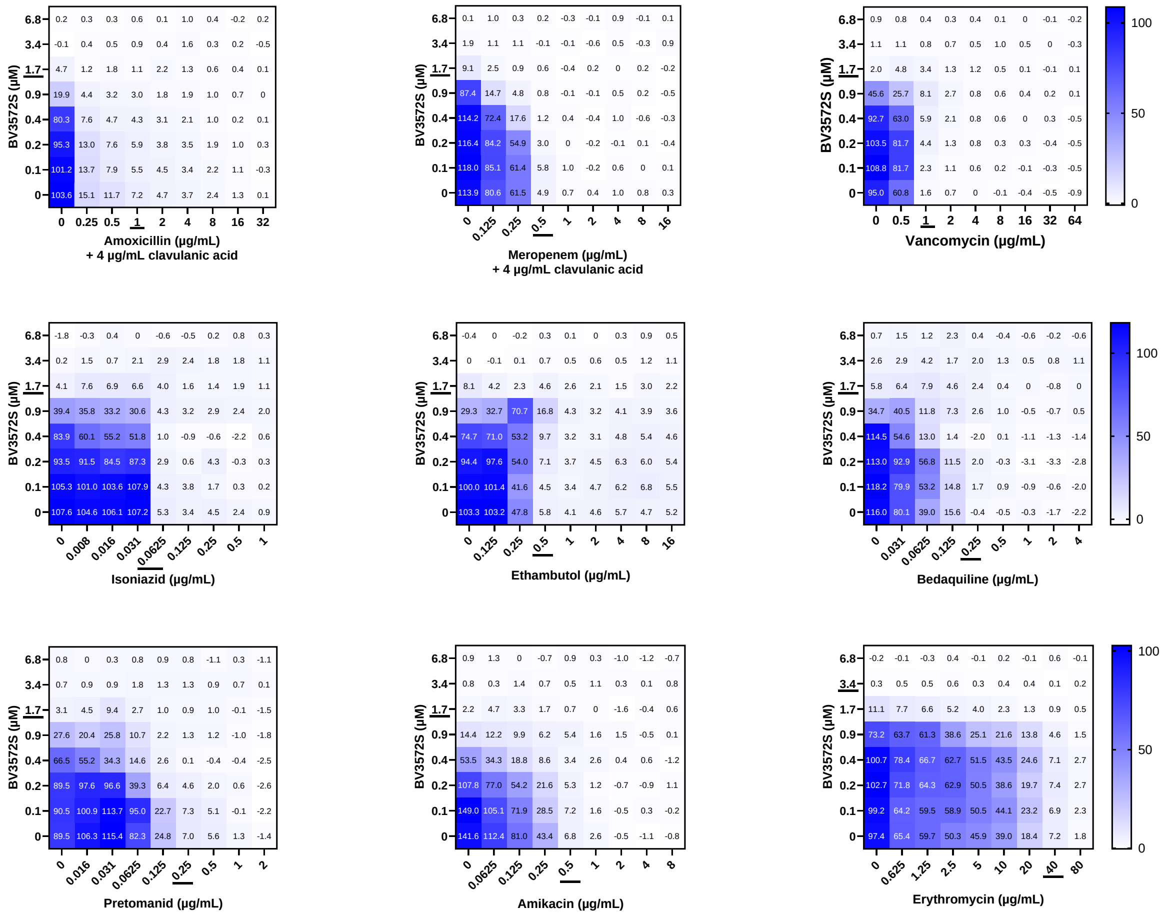

**Supplementary Fig. 3.** Heatmap depictions of normalized fluorescence of *Mtb* H37Rv treated with combinations of BVL3572S and the indicated antibiotics. Underlined values indicate the $MIC_{90}$ of each antibiotic.

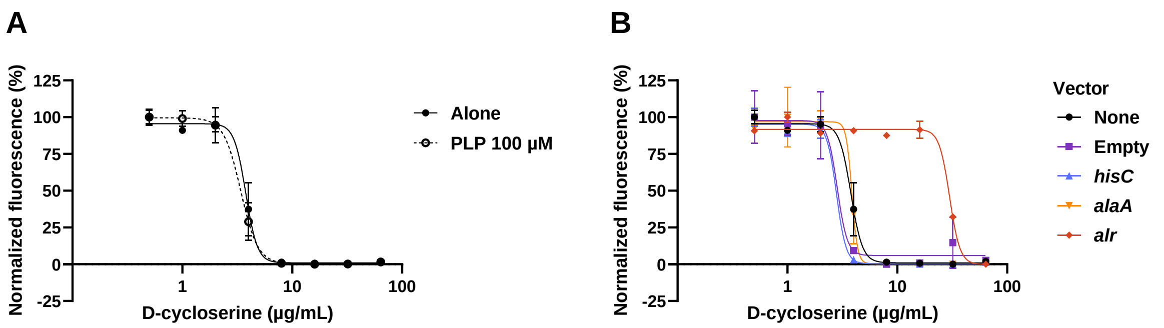


**Supplementary Fig. 4. (A)** Effect of PLP supplementation on the growth inhibitory activity of D-cycloserine against *Mtb*. **(B)** Effect of *hisC*, *alaA*, or *alr* overexpression on the growth inhibitory activity of D-cycloserine compared to *Mtb* alone or containing the empty pMV261 vector. Data shown are means ± standard deviation of two technical replicates and are representative of three biological replicates.
